# Expanded Genome and Proteome Reallocation in a Novel, Robust *Bacillus coagulans* Capable of Utilizing Pentose and Hexose Sugars

**DOI:** 10.1101/2024.07.15.603586

**Authors:** David Dooley, Seunghyun Ryu, Richard J. Giannone, Jackson Edwards, Bruce S. Dien, Patricia J. Slininger, Cong T. Trinh

## Abstract

*Bacillus coagulans*, a Gram-positive thermophilic bacterium, is recognized for its probiotic properties and recent development as a microbial cell factory. Despite its importance for biotechnological applications, the current understanding of *B. coagulans*’ robustness is limited, especially for undomesticated strains. To fill this knowledge gap, we characterized the metabolic capability and performed functional genomics and systems analysis of a novel, robust strain, *B. coagulans* B-768. Genome sequencing revealed that B-768 has the largest *B. coagulans* genome known to date (3.94 Mbp), about 0.63 Mbp larger than the average genome of sequenced *B. coagulans* strains, with expanded carbohydrate metabolism and mobilome. Functional genomics identified a well-equipped genetic portfolio for utilizing a wide range of C5 (xylose, arabinose), C6 (glucose, mannose, galactose), and C12 (cellobiose) sugars present in biomass hydrolysates, which was validated experimentally. For growth on individual xylose and glucose, the dominant sugars in biomass hydrolysates, B-768 exhibited distinct phenotypes and proteome profiles. Faster growth and glucose uptake rates resulted in lactate overflow metabolism, which makes *B. coagulans* a lactate overproducer; however, slower growth and xylose uptake diminished overflow metabolism due to the high energy demand for sugar assimilation. Carbohydrate Transport and Metabolism (COG-G), Translation (COG-J), and Energy Conversion and Production (COG-C) made up 60–65% of the measured proteomes but were allocated differently when growing on xylose and glucose. The trade-off in proteome reallocation, with high investment in COG-C over COG-G, explains the xylose growth phenotype with significant upregulation of xylose metabolism, pyruvate metabolism, and TCA cycle. Strain B-768 tolerates and effectively utilizes inhibitory biomass hydrolysates containing mixed sugars and exhibits hierarchical sugar utilization with glucose as the preferential substrate.

**IMPORTANCE:** The robustness of B. coagulans makes it a valuable microorganism for biotechnology applications, yet this phenotype is not well understood at the cellular level. Through phenotypic characterization and systems analysis, this study elucidates the functional genomics and robustness of a novel, undomesticated strain, B. coagulans B-768, capable of utilizing inhibitory switchgrass biomass hydrolysates. The genome of B-768, enriched with carbohydrate metabolism genes, demonstrates high regulatory capacity. The coordination of proteome reallocation in Carbohydrate Transport and Metabolism (COG-G), Translation (COG-J), and Energy Conversion and Production (COG-C) is critical for effective cell growth, sugar utilization, and lactate production via overflow metabolism. Overall, B-768 is a novel, robust, and promising B. coagulans strain that can be harnessed as a microbial biomanufacturing platform to produce chemicals and fuels from biomass hydrolysates.

## INTRODUCTION

Lignocellulosic biomass derived from bioenergy crops (e.g., poplar, switchgrass) or agricultural residues (e.g., corn stover) are renewable and sustainable feedstocks(1, 2). These feedstocks can be produced at a scale of 1 billion tons per year in the US, reducing reliance on petroleum to make fuels, chemicals, and materials(3). Lignocellulosic biomass is comprised of cellulose, hemicellulose, and lignin(4). These polymers can be isolated to make consumables(5, 6), advanced materials(7), and other valuable commodities. Alternatively, these polymers can be further broken down to monomers through chemical pretreatment followed by enzymatic saccharification to generate biomass hydrolysates that consist of pentose (C5) and hexose (C6) sugars and byproducts (e.g., organic acids, furans, and phenolics). Through fermentation, microbial cell factories can be harnessed as biocatalysts to convert these sugars into various classes of fuels, chemicals, and materials(2, 8, 9) including carboxylic acids(10), alcohols(11–13), aldehydes(14), ketones(15), amines(16), esters(17, 18), polyhydroxyalkanoates(19), and protein-based polymers(20). Biomanufacturing is thus important for a circular bioeconomy where renewable and sustainable feedstocks can be effectively utilized to make renewable goods(21).

One technical barrier to biomanufacturing is creating robust microbial catalysts capable of transforming biomass hydrolysates into fuels, chemicals, and materials. Robustness is defined as the efficacy by which microbes can convert biomass hydrolysates into a target chemical under harsh conditions that cause biocatalyst poisoning(22, 23). Robustness is a complex phenotype that requires controlled expression of multiple genes. For instance, the effective assimilation of C5 and C6 sugars in biomass hydrolysates is a complex phenotype, requiring a highly regulated gene network(24). Coping with chemical inhibitors such as pretreatment byproducts in biomass hydrolysates and target products generated during fermentation forces cells to optimize their cellular resources at the systems level(25). Therefore, deploying robust microbial catalysts, whether natural isolates or engineered microorganisms, for efficient conversion of lignocellulosic biomass to a target product is critical for biomanufacturing.

The robustness and unique metabolic capability of *Bacillus coagulans* make it a promising biomanufacturing platform. *B. coagulans* (also known as *Weizmannia*/*Heyndrickxia coagulans*) is a GRAS (generally recognized as safe) Gram-positive bacterium and facultative thermophilic anaerobe. *B. coagulans* grows on a variety of carbohydrates(26–28) at a wide range of temperatures(29) and pHs(30) and is capable of overproducing lactate(31), thermostable enzymes(31), and antimicrobial peptides(31–33). Due to its unique metabolic capability and inherent robustness, *B. coagulans* is also utilized for probiotic formulations(34). Even though it is a promising biomanufacturing platform, the fundamental understanding of its robustness and underlying genetics is limited.

In this study, we characterized the robustness physiology of *B. coagulans* B-768, a novel, undomesticated strain. We first assessed its unique metabolic capability by growing it on various C5 (arabinose, xylose), C6 (glucose, galactose, mannose), and C12 (cellobiose) sugars at a wide range of temperatures and evaluated its tolerance to high lactate concentrations. Genome sequencing and annotation were performed to provide insights into its observed metabolic capability. Subsequent characterization of B-768 growing on xylose, glucose, a mixture of xylose and glucose, and biomass hydrolysate, coupled with quantitative proteomics, validated the functions of genes involved in sugar utilization and elucidated how cells dynamically reallocate their proteomes in response to different carbon sources. Finally, we examined the regulation of sugar utilization in B-768 and uncovered the cellular processes and molecular drivers responsible for B-768’s robust growth on undetoxified hydrolysates.

## RESULTS AND DISCUSSION

### *B. coagulans* B-768 is a robust biomanufacturing platform microorganism

Utilization of a wide range of biomass-derived sugars is crucial for a robust biomanufacturing platform microorganism. Strain B-768 was grown on glucose, xylose, and cellobiose -the dominant sugars present in biomass hydrolysates -at a range of temperatures (30– 59 °C, Figs. 1A–C). Optimum cell growth was observed between 45–53 °C with a maximum growth rate of 0.4 h^-1^ on glucose and 0.2 h^-1^ on xylose or cellobiose. Next, B-768 was grown in media containing galactose, mannose, or arabinose at 50 °C in addition to the previous sugars (Fig. 1D). Glucose and mannose were the most efficiently used carbon sources, followed by galactose, xylose, arabinose, and cellobiose, which all yielded similar maximum specific growth rates and maximum cell biomasses (Figs. 1D, S1A). In addition to the assimilation efficiencies of various carbon sources, tolerance to high product concentrations is critical to determine cellular robustness. Since *B. coagulans* is well-known for its high lactate production, we evaluated the growth of B-768 in cultures containing various lactate concentrations with glucose as a sole carbon source (Figs. 1E, S1B). The maximum specific growth rate was reduced as lactate concentration was increased, and B-768 could tolerate up to at least 30 g/L lactate. Since this experiment was performed in a pH-uncontrolled environment, it was hypothesized that higher lactate tolerance could be achieved when the medium pH was maintained at an optimum level. To evaluate this hypothesis, a pH-controlled fermentation was performed with 80 g/L glucose as the sole carbon source. As expected, when the pH was maintained at 6.5, B-768 produced up to 60 g/L lactate at 92.7% of the theoretical maximum yield (Fig. 1F). These results highlight the robustness of B-768 for metabolizing biomass-derived sugars, growing over a broad range of temperatures, and selectively producing lactic acid at high yields.

**Figure 1:**
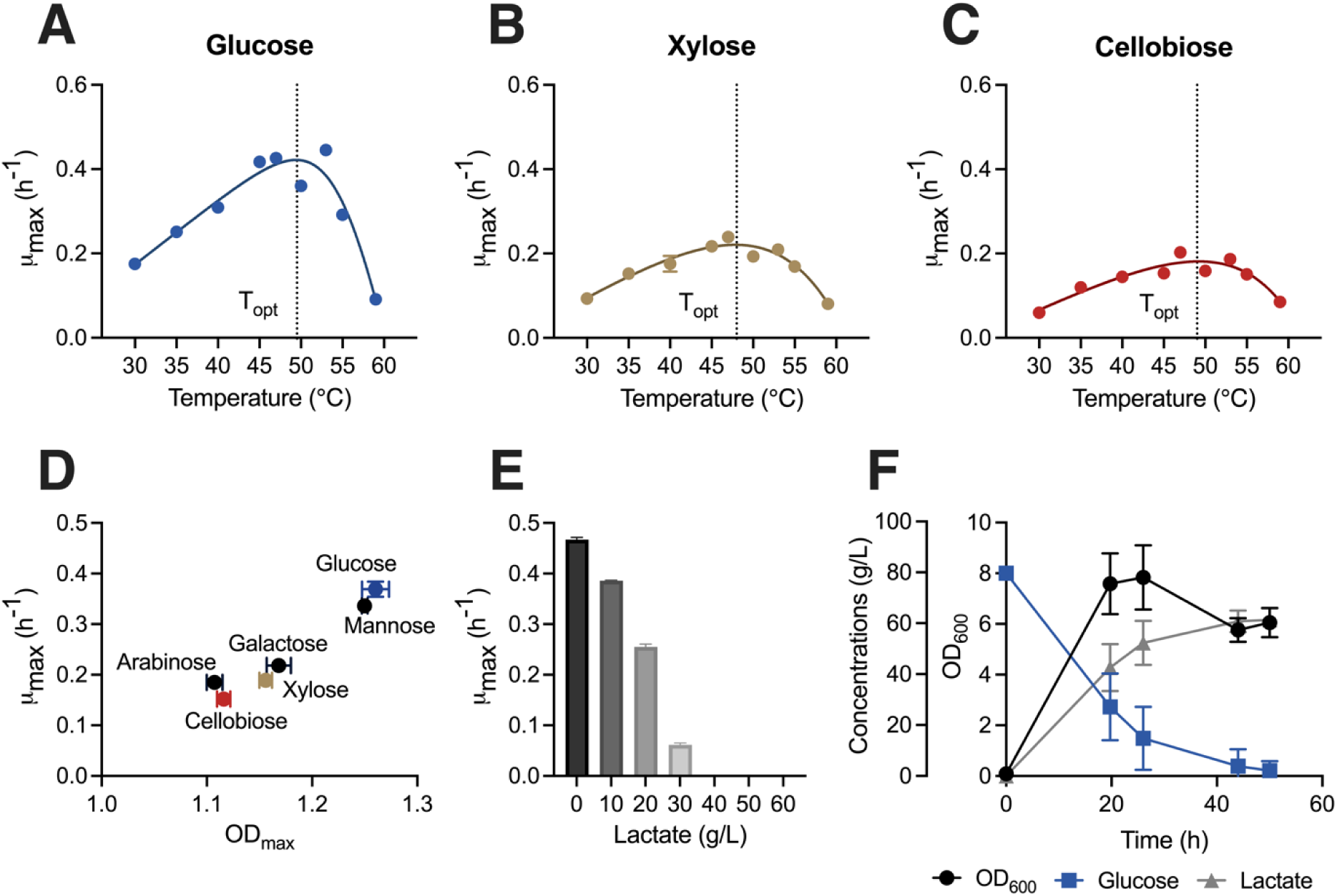
Physiological characterization of *B. coagulans* B-768 reveals cellular robustness and efficient utilization of various carbon sources at various temperatures. (A–C) Dependence of specific growth rates of B-768 on temperatures when grown on (A) glucose, (B) xylose, and (C) cellobiose. (D) Carrying capacity and maximum specific growth rate of B-768 growing on various C5, C6, and C12 fermentable sugars at 50 °C. (E) Effect of lactate concentrations on maximum specific growth rates of B-768 growing on glucose at 50 °C. (F) Kinetic profiles of cell growth, glucose consumption, and lactate production in a pH-controlled bioreactor at 50 °C. Data points represent 1 mean ± 1 standard deviation of three biological replicates.

### Functional and comparative genomics of *B. coagulans* B-768

#### Genome sequence of B-768 and its relatedness to other B. coagulans strains

The genome of B-768 was sequenced in an attempt to elicit the genetic basis of its physiological robustness. The genome assembly size of B-768 is 3,944,476 base pairs long with 45.7% GC content, which is 630 Kbp larger than the average *B. coagulans* genome and is the largest reported genome for *B. coagulans* (Fig. 2). Strain B-768 contains 4,059 genomic features, of which 3,863 (95.2%) are coding sequences (CDS). Phylogenomic analysis of a 55-strain *B. coagulans* pan-genome compared to model strains 36D1 and DSM1 showed that B-768 belongs to the same clade as 36D1 (clade A) and is radically diverged from DSM1 (clade B, Fig. 3A). Also, its genome is 11% larger than 36D1 and 17% larger than DSM1 (Table 1). Concordantly, there are ∼20% more CDS in B-768 compared to 36D1 or DSM1. Strains B-768 and 36D1, but not DSM1, are capable of metabolizing C5 sugars (e.g., xylose and arabinose)(31, 35–37). Other strains reported to metabolize xylose, including NL01(27, 38) and DSM 2314(39), also belong to clade A. The ability to efficiently use C5 sugars could aid strains in occupying an ecological niche and incentivize the acquisition of relevant genomic material through vertical and horizontal gene transfers. This hypothesis is supported by the fact that genomes from clade A are on average ∼171 Kbp larger than those for clade B. Though further characterization of xylose and arabinose usage in *B. coagulans* strains is needed to confirm this hypothesis, C5 metabolism could partially explain the genetic differences underlying the topology of the *B. coagulans* pan-genome.

**Figure 2:**
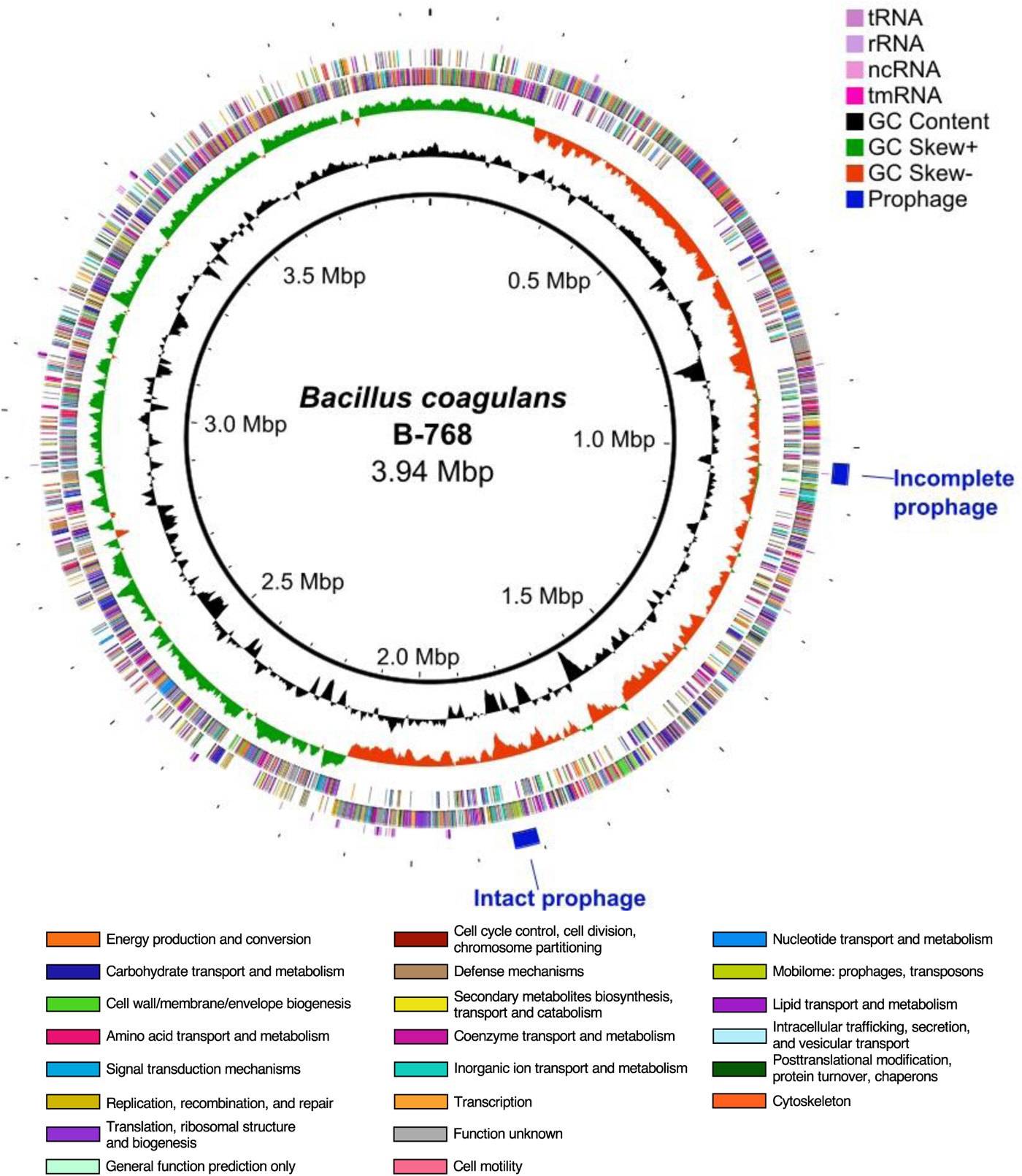
Circular genome map of *Bacillus coagulans* B-768. Tracks from outside to in are as follows: PHASTEST 3.0 prophage regions, non-coding RNAs, forward and reverse strand CDS colored by COG category, GC Skew = (G - C)/(G + C), and GC content. Abbreviations: tRNA, transfer RNA; rRNA, ribosomal RNA; ncRNA, non-coding RNA; tmRNA, transfer-messenger RNA.

**Figure 3:**
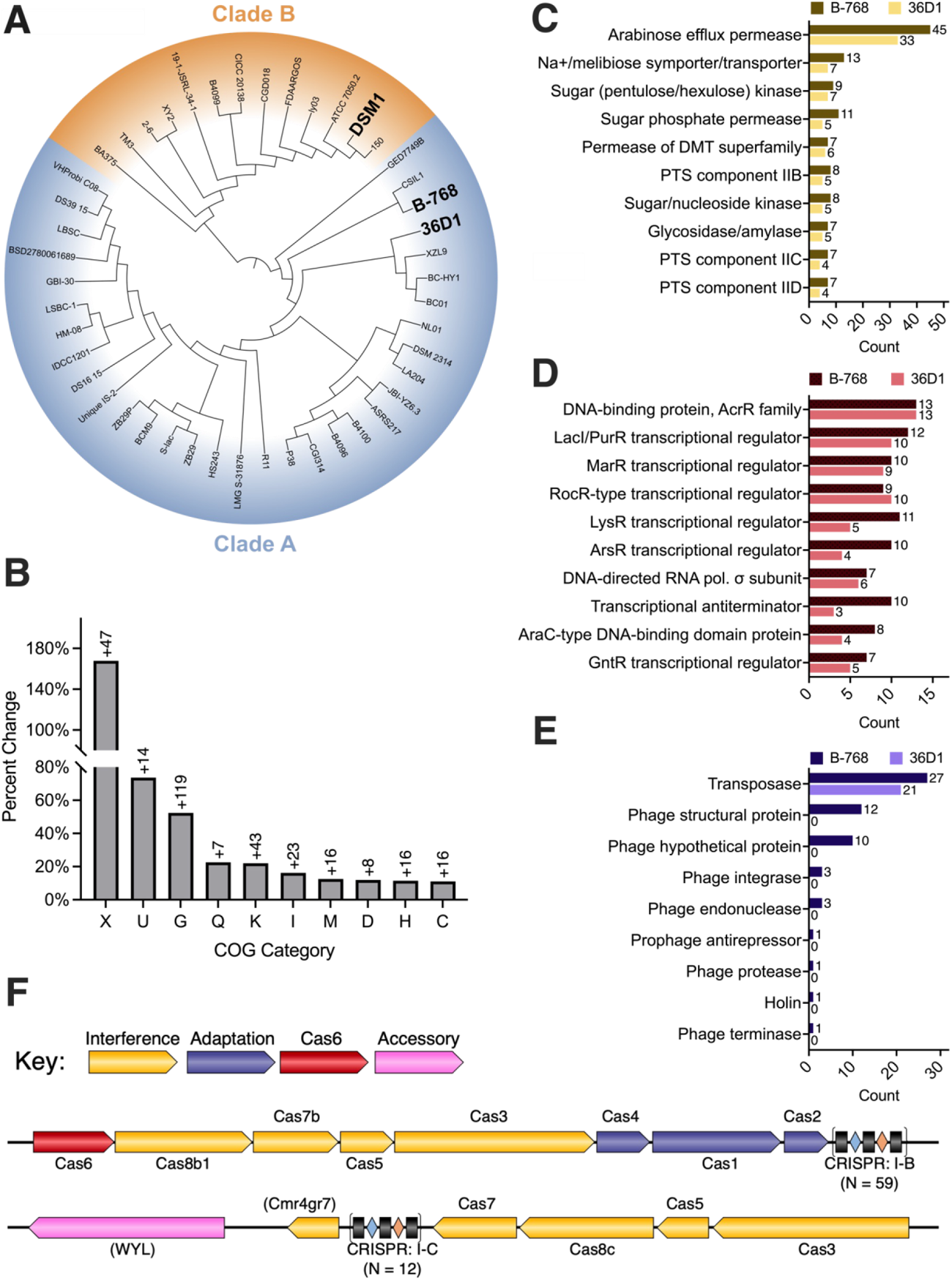
Comparative and functional genomics of *B. coagulans* B-768. (A) Core genome alignment of *B. coagulans* pan-genome (n = 55). (B) Percent change of COG categories between strains B-768 and 36D1. Numbers above the bars indicate the excess number of proteins in B-768 relative to 36D1. Full COG Category names are listed at the end of the figure caption. (C–E) Breakdown of most abundant proteins in (C) Carbohydrate Metabolism and Transport, (D) Transcription, and (E) Mobilome COG categories. (F) Predicted CRISPR-Cas operons of B-768. COG Categories: U = Intracellular Trafficking, Secretion, and Vesicular Transport; X = Mobilome; G = Carbohydrate Transport and Metabolism; Q = Secondary Metabolites Biosynthesis, Transport and Catabolism; K = Transcription; I = Lipid Transport and Metabolism; M = Cell Wall/Membrane/Envelope Biogenesis; D = Cell Cycle Control, Cell Division, Chromosome Partitioning; H = Coenzyme Transport and Metabolism; C = Energy Production and Conversion.

**Table 1:**
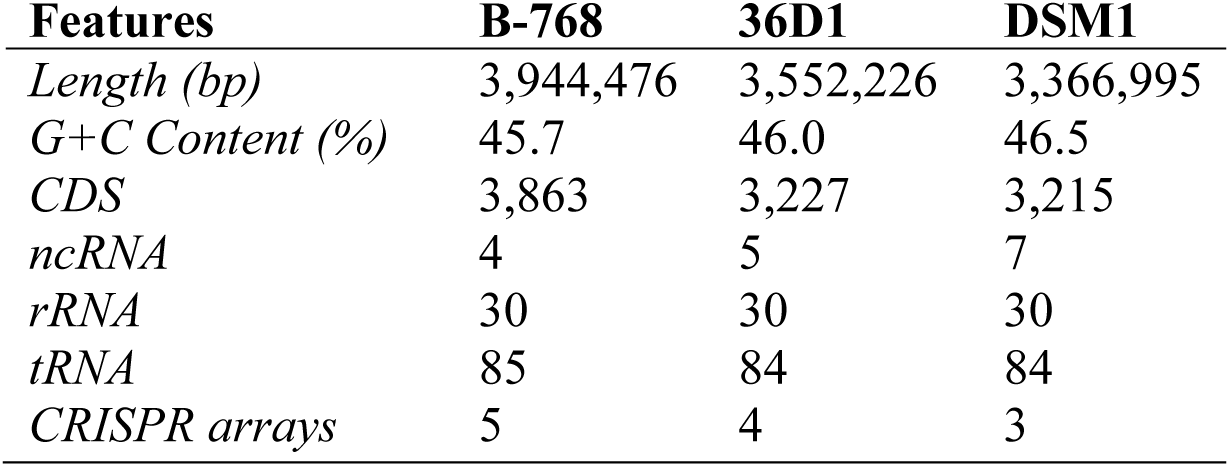
Summary of *B. coagulans* B-768 assembly and comparison with reference genomes.

#### The larger genome of B-768 provides potential gains-of-function for enhanced cellular robustness

To better understand the biological functions that may be associated1 with the added genomic material of B-768, we conducted a comparative COG (Clusters of Orthologous Genes) analysis by calculating the percent change of CDS belonging to each COG category between B-768 and its closest reference, 36D1 (Fig. 3B–E). The percent change was calculated as follows: 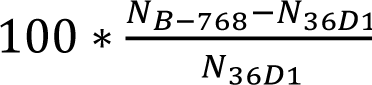, where N is the number of proteins belonging to the specified COG. Several putative phage proteins found using PHASTEST(40) not recognized by automated COG annotation(41) were manually added to the Mobilome COG. The percent change of the top 10 COGs is shown in Fig. 3B. B-768 contains more genes than 36D1 for nearly every COG category with several categories being heavily expanded, such as the Mobilome (COG-X), Intracellular Trafficking, Secretion, and Vesicular Transport (COG-U), Carbohydrate transport and metabolism (COG-G), and Transcription (COG-K, Fig. 3B). Detailed examination of the individual proteins involved in these COGs can shed light into the robustness of B-768, specifically COG-G and COG-K due to their relevance to carbon utilization (Fig. 3C & 3D) and COG-X due to its role in horizontal gene transfer and genome composition (Fig. 3E).

#### Unique carbohydrate metabolism, transport, and regulation of B-768

Carbohydrate Metabolism and Transport is the largest functional category in B-768, underlying the observed robust carbon metabolism. B-768 contains 52.4% more proteins in COG-G than 36D1, and more proteins were identified for all protein classes (Fig. 3C). Top protein classes include arabinose efflux permeases, Na+/melibiose symporters/transporters, sugar kinases, and sugar permeases. The overrepresentation of transporters among this group suggests B-768’s vigorous carbon metabolism is partly due to improved uptake of carbon substrates. Transcriptional regulators are known to play key roles in regulating the use of carbon sources. A breakdown of the COG-K reveals B-768 has at least 16 more transcriptional regulators than 36D1, belonging to the LacI/PurR, LysR, ArsR, and GntR families (Fig. 3D). LacI and PurR regulators are mostly allosteric repressors of carbon and nucleotide synthesis, respectively, and may control one (e.g. LacI) or many (e.g. PurR) operons(42). Given the ability of B-768 to utilize various carbon sources (Fig. 1D), an enlarged complement of LacI/PurR regulators likely helps coordinate the use of available sugars. LysR and GntR regulators act globally as either activators or repressors of various cellular processes including, but not limited to, metabolism, virulence, motility, and intercellular communication(43, 44). ArsR regulators repress operons linked to metal stress under normal physiological conditions. Upon binding with multivalent metal or metalloid ions, the associated operons are derepressed and can provide resistance to otherwise toxic concentrations of metals/metalloids(45). An excess of 6 ArsR regulators in B-768 may enable it to survive in metal-rich environments where other *B. coagulans* strains cannot thrive.

#### Mobilome of B-768 reveals its capability to expand its genome

B-768 contains 168% more proteins in its Mobilome—which includes prophages, transposons, and plasmid mobilization and maintenance machinery—compared to 36D1 (Fig. 3E). Six more transposases, which are capable of rearranging, adding, or removing genomic material, were identified from B-768’s chromosome. The most significant portion of B-768’s expanded Mobilome, however, is attributed to two putative prophage regions that encode 32 CDS. In contrast, 36D1 has no prophages or phage-related proteins. Prophage-like entities are commonly found in probiotic strains and can provide resistance to similar phages(46, 47). The first prophage region is 32.7 kb long (genomic coordinates: 1024169–1056950) and is predicted to be incomplete due to the absence of various necessary phage genes such as portal, capsid, terminase, and tail genes. It does, however, encode for 2 integrases, 1 protease, 1 endonuclease, 1 antirepressor, and several other unannotated phage-like proteins which may contribute to bacterial fitness and/or horizontal gene transfer. The second prophage is 38.3 kb long (genomic coordinates: 1805785–1844080) and is putatively intact, possessing all major proteins required for phage infection. Many of the closest homologs for B-768’s putative phage proteins are from *Bacillus* phages, but several come from *Corynebacterium*, *Agrobacterium*, and *Enterococcus* phages as well, meaning the exact origin and classification of the prophage regions remain unclear. The existence of prophages in *B. coagulans* strains has been previously reported, but in all cases the phages were deemed non-functional(48, 49). Given the well-established role of bacteriophages in horizontal gene transfer, the presence of two putative prophages could help explain the origin of B-768’s expanded genetic repertoire. An operational phage in *B. coagulans* could potentially be useful for introducing DNA via transduction, enabling various genetic engineering approaches. Together, these findings provide a glimpse into the expanded genome of B-768 that facilitates its extraordinary cellular robustness.

#### Defense systems in B-768: restriction-modification and CRISPR-Cas

Bacteriophage contamination is a common problem in bacterial fermentation and can lower product yields and either slow or halt fermentnations(50). Bacteria have developed numerous defense systems for rebuffing bacteriophage infection, including innate restriction-modification (R-M) and adaptive CRISPR (clustered regularly interspaced short palindromic repeats)-Cas (CRISPR-associated). To elucidate the robustness of B-768, we evaluated its innate and adaptive defense systems to assess its potential resistance to phage infection and suitability for industrial application.

Restriction-modification systems recognize and cut foreign DNAs having different methylation patterns than the host. R-M systems are categorized into four types, Type I–IV, based on genetic structure and biochemical function. Type I systems are multi-subunit complexes composed of the R (restriction endonuclease), M (methyltransferase), and S (DNA-specificity) subunits expressed from a single operon. Type II and III systems are comprised of restriction endonuclease and methyltransferase subunits, but while Type II R and M proteins function independently, Type III restriction enzyme activity requires an R-M complex. Type IV systems are effectors that may be encoded by either 1 or 2 genes(51). Type I, II, and III restriction endonucleases recognize and cleave unmethylated DNA at a specific DNA motif, while Type IV restriction endonucleases cleave DNA that is methylated differently from the host. Our genomic analysis shows that B-768 has 11 predicted R-M proteins including 7 Type I, 2 Type II, and 2 Type IV proteins (Table 2). No Type III R-M proteins were found. In B-768, a complete Type I R-M system is encoded within a single operon spanning from 2754339–2761240. We found that B-768 also has 4 additional Type I R submodules co-localized from 3201231–3202333, but it is unclear whether these isolated proteins participate in Type I R-M activity in B-768. Both Type II R-M proteins in B-768 are contained in an operon from 2170437–2176691. B-768’s Type IV genes are co-localized, meaning they likely encode for a 2-subunit effector. In total, B-768 has 3 distinct and intact R-M systems that enable robust restriction of foreign DNAs.

**Table 2:**
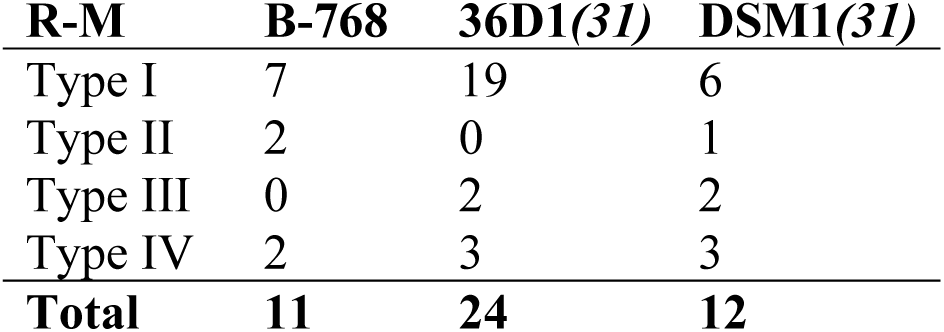
Number of genes in restriction-modification (RM) systems.

CRISPR-Cas systems are functionally distinct immune systems found in archaea, bacteria, and even huge bacteriophages(52–54). These systems work in a two-step process: adaptation and interference. In adaptation, stretches of foreign DNA (protospacers) are integrated into CRISPR arrays within the host genome and stored for future use. These short fragments are called spacers. During interference, spacers are transcribed, form a complex with Cas effector proteins, and guide a ribonuclease to a cognate protospacer, where the target DNA is cleaved. Our genomic analysis found that B-768 possesses 5 putative CRISPR arrays containing 145 spacers. Three CRISPR arrays have no associated Cas proteins near them and 2 have accompanying Type I-B and I-C CRISPR machinery (Fig. 3F). The Type I-B CRISPR system is predicted to have complete interference and adaptation arsenals, while the Type I-C system appears to only possess interference-related genes. Analysis of B-768’s entire set of spacers reveals CRISPR RNAs targeting both phages and plasmids, which indicates B-768 has a history of deploying CRISPR-Cas against foreign invaders of various origins (Table 3). In addition to robust and programmable defense against deleterious phages, B-768’s endogenous CRISPR systems could hold promise for editing *B. coagulans* and other industrially relevant thermophiles.

**Table 3:**
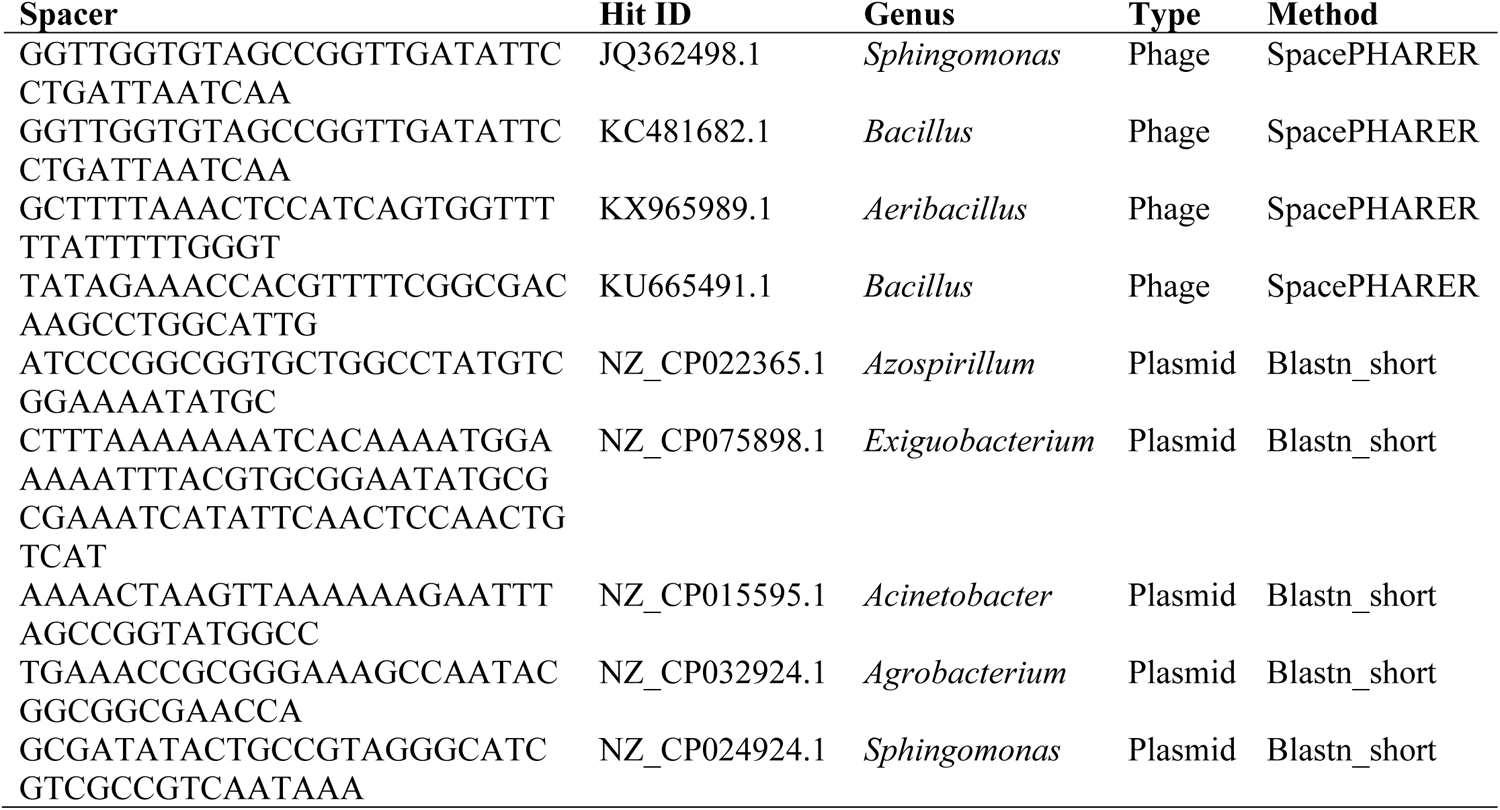
Identification of spacers in B-768 CRISPR arrays.

#### Lignocellulosic biomass-derived sugar metabolism in B-768

The C6 sugars (e.g., glucose) are mainly transported inside the cells using the phosphoenolpyruvate (PEP):carbohydrate phosphotransferase system (PTS) (Dataset 1). The phosphorylated sugars (e.g., glucose-6-phosphate) are assimilated via the Embden-Meyerhof-Parnas (EMP) and/or Entner-Doudoroff (ED) pathways. We identified the full set of necessary genes for the EMP pathway in B-768 (Fig. 4, Dataset 2). On the other hand, B-768 contains an incomplete ED pathway due to the absence of 6-phosphogluconate dehydratase (EDD), which converts 6-P-gluconate into 2-keto-3-deoxy-6-phosphogluconate (Fig. 4, Dataset 2). B-768 does, however, possess 6-phosphogluconate dehydrogenase (6PGD) that converts 6-phosphogluconate into ribulose-5-phosphate, one of the end products in the pentose phosphate pathway. In summary, glycolysis is the primary pathway for C6 sugar degradation in B-768, with possible metabolic flux through a hybrid ED and pentose phosphate pathway route.

**Figure 4:**
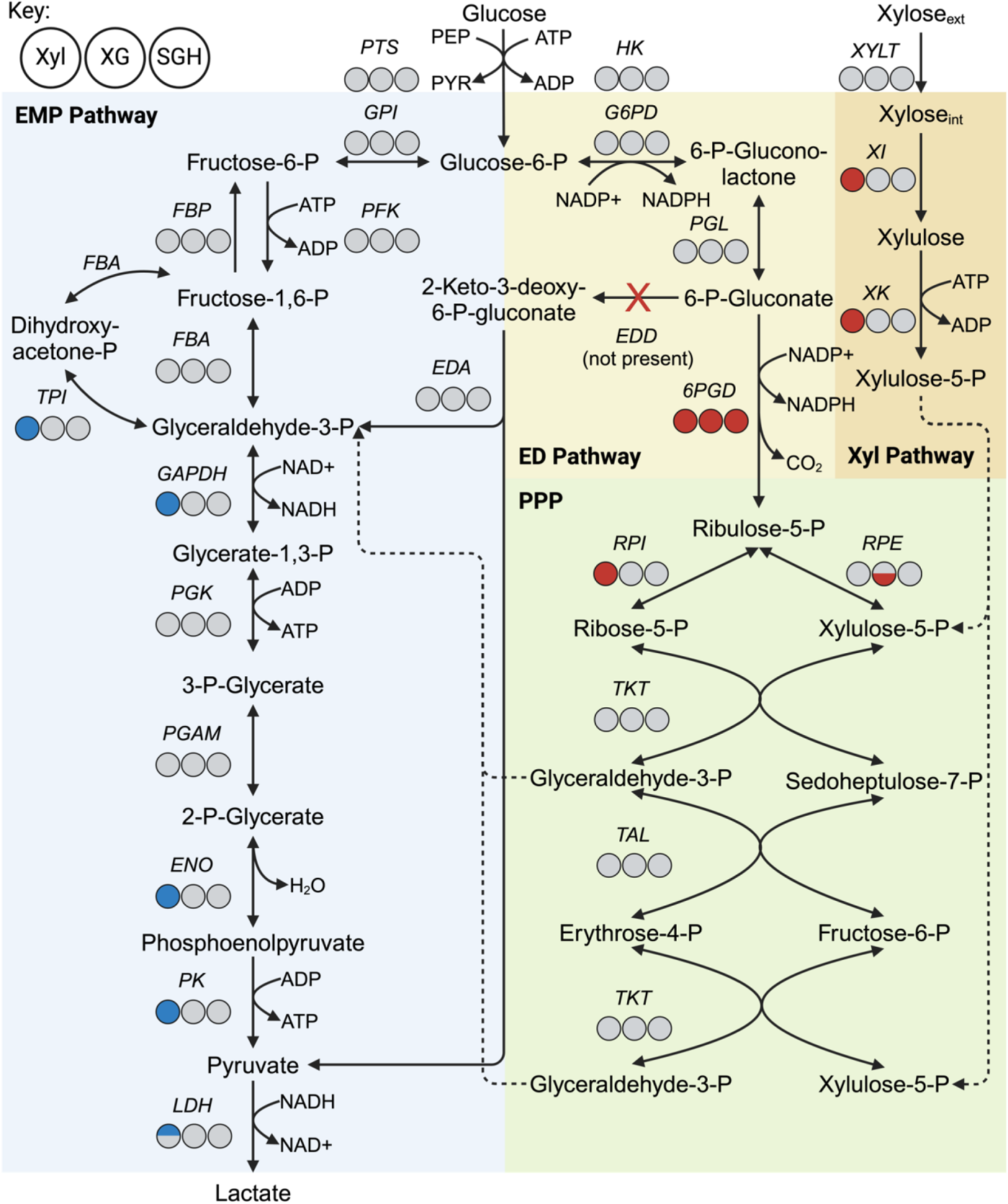
Central carbon metabolism of B-768. Circles from left to right represent the differential expression of a given protein in 1.) xylose vs. glucose, 2.) xylose and glucose mixture vs. glucose, and 3.) switchgrass hydrolysate vs. glucose conditions. Red = upregulated with log_2_-fold change ≥ 1 and q < 0.05; Blue = downregulated with log_2_-fold change ≤ -1 and q < 0.05; Grey = Not Significant. Paralogs of the same enzyme were plotted as half circles with the most abundant paralog on top.

Since B-768 grows on xylose (Figs. 1B, 1D), it is expected that B-768 contains xylose assimilation genes. Transport is the first step of carbon utilization and B-768 contains two putative xylose transporters, xylose H+-symporter (XYLT, encoded by *xylT*) and an ABC-type xylose transporter (*xylFGH*). These were identified using 36D1 xylose transporters as reference genes (Dataset 1). Xylose transported inside of the cell via xylose transporters is metabolized by two key enzymes: xylose isomerase (XI, encoded by *xylA*), which converts D-xylose into D-xylulose, and xylulokinase (XK, encoded by *xylB*), which converts D-xylulose into D-xylulose-5-phosphate. Typically, *xylA*, *xylB*, and *xylT* are encoded alongside *xylR*—a transcriptional repressor of the *xyl* operon(55, 56)— in a single operon, but in B-768 *xylT* is located separately over 1 Mbp away (Fig. 4, Dataset 2). Once xylose is converted into D-xylulose-5-phosphate, it can be further assimilated by either the pentose phosphate pathway (PPP) or the phosphoketolase pathway (PKP). The PKP results in equimolar production of lactate and acetate in lactic acid bacteria(57), while the PPP yields homolactic acid, making the PPP desirable for lactate production from pentoses(36). B-768 contains a full complement of PPP genes but lacks the critical PKP enzyme, phosphoketolase (Dataset 2). Given the lack of phosphoketolase, B-768 mainly utilizes the homolactic acid PPP to degrade xylose, setting it apart from most other lactic acid bacteria that prefer the PKP(58). This is an essential trait for ensuring high lactic acid yields on cellulosic sugar mixtures.

B-768 also contains genes that putatively allow it to use xylosides. Xylosides are released when xylan or xyloglucan are depolymerized and may be converted into D-xylose by ⍺-or β-xylosidases(59). Putative xyloside symporter (*xynT*) and ⍺-xylosidase (*xylS*) genes were found in the *xyl* operon in B-768, and three β-xylosidases were scattered throughout the rest of the genome. While closely related strain 36D1 has a copy of *xynT*, it does not possess any known xylosidases(31). To determine the origin of the four xylosidase proteins, BLASTp analysis was performed against the non-redundant protein database (Table 4). This search revealed homologs in closely related species for all but one of the xylosidases. Enzyme 4-beta-xylosidase located from 665440–666999 had top hits primarily belonging to *Enterococcus* spp., with the top hit occurring in *Melissococcus plutonius*. This provides another example of *B. coagulans* B-768 utilizing horizontal gene transfer to expand its carbon-utilizing potential. This is useful trait because xylan oligomers are competitive inhibitors of cellulases.

**Table 4:**
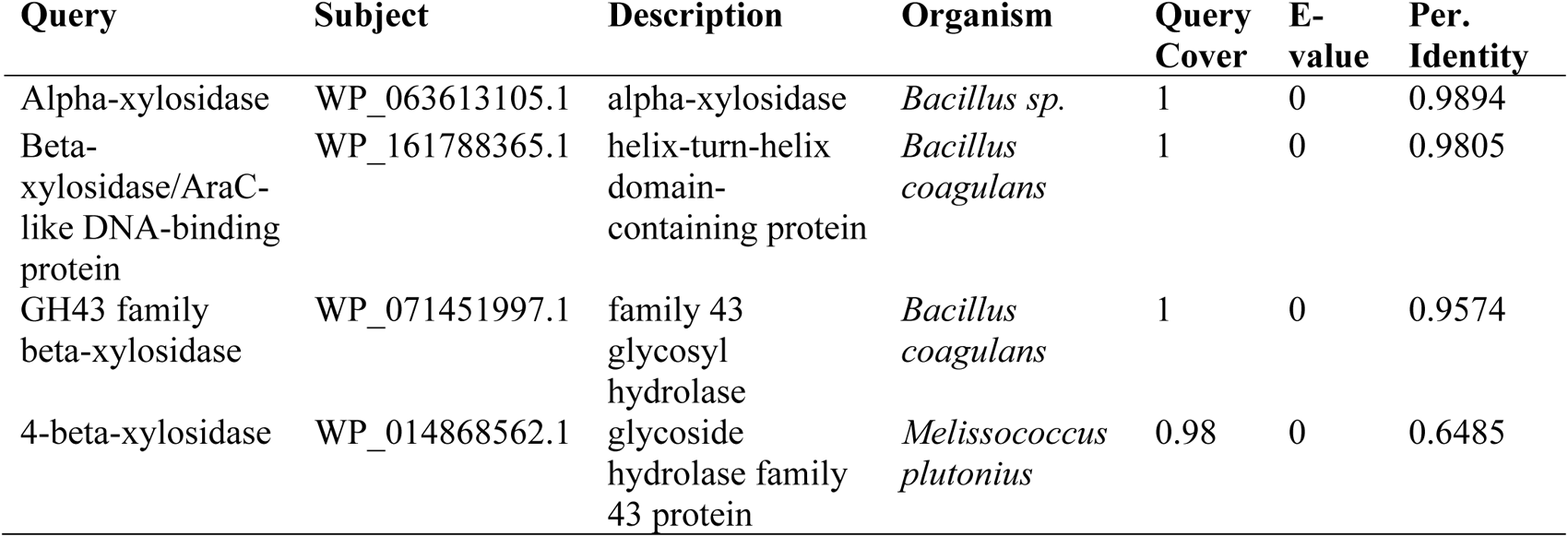
BLASTp top hits for B-768 xylosidase proteins.

Cellobiose assimilation is key to bypassing carbon catabolite repression (CCR), wherein xylose assimilation is inhibited by glucose. Cellobiose utilization in *B. coagulans* is mainly mediated by the five-member CELO1 operon encoding three phosphotransferase system (PTS) subunits EIIA (*celA*), EIIB (*celB*), EIIC (*celC),* a 6-phospho-β-glucosidase (*pbgl*), and a transcriptional regulator (*celR*)(27). Some strains like *B. coagulans* NL01, 36D1, 2-6, and HM08 have an additional cellobiose operon, CELO2, which encodes paralogs of *celABCR* along with an unknown protein (*celX*)(27). Using 36D1 as a reference strain, we identified cellobiose assimilation pathway genes in B-768. In B-768, both CELO1 and CELO2 are fully intact and located from 1361260–1367923 and 1464816–1469898, respectively. Notably, B-768 contains two additional copies of 6-phospho-β-glucosidase—the enzyme catalyzing the rate-limiting step in the breakdown of cellobiose to glucose(60, 61)—which may improve cellobiose catabolism.

Due to B-768’s metabolic capability to assimilate galactose, mannose, and arabinose (Fig. 1D), we also investigated the catabolic pathways of these monomeric sugars. Galactose metabolism in bacteria is mediated by the ubiquitous four-enzyme Leloir pathway, containing the enzymes galactose mutarotase, galactokinase, galactose-1-phosphate uridylyltransferase, and UDP-galactose 4-epimerase(62). B-768 possesses all four genes in an operon from 1161267– 1165598 with an additional copy of galactose mutarotase (Dataset 1). Mannose metabolism in *Bacillus subtilis* is mediated by a mannose transporter (*manP*), a mannose-6-phosphate isomerase (*manA*), and a mannose activator (*manR*)(63). In *B. coagulans*, mannose metabolism is much the same, except the mannose transporter is mediated by the mannose PTS system encoded by 4 subunit genes (*manEIIABCD)* instead of a single gene (Dataset 1). B-768 possesses homologs for these genes, but *manA* is absent from the operon containing *manR* and *manEIIABCD* (Dataset 1). All canonical arabinose utilization proteins are present in B-768, many of which possess several putative paralogs (Dataset 1). Most of these paralogs are either for arabinose/arabinan permeases or transcriptional regulators of the *ara* regulon. This is consistent with the finding that most of B-768’s genes involved in carbon metabolism putatively encode for C5/C6 sugar kinases, permeases, and transcriptional regulators (Fig. 3C). Overall, B-768 has accrued an extensive array of genes dedicated to metabolizing a wide variety of sugars.

### Distinct phenotypes and proteomes of B-768 for utilization of biomass-derived sugars

#### B-768 grew faster on glucose than xylose, resulting in lactate overflow metabolism

Since xylose and glucose are the most dominant C5 and C6 sugars in biomass hydrolysates, we first aimed to elucidate the phenotypes and metabolic responses of B-768 growing on these sugars. When supplied with 10 g/L glucose, B-768 grew at a rate of 0.59 ± 0.01 h^-1^. Within 8 h, it achieved a maximum growth of 3.45 ± 0.11 OD_600_ and exhausted all the glucose with a glucose uptake rate of 1.67 ± 0.15 g glucose/OD_600_/h (Fig. 5A, 5B). B-768 produced lactate as the major fermentative product at a titer of 8.72 ± 0.05 g lactate/L, a productivity of 1.46 ± 0.03 g lactate/OD_600_/h, and a yield of 0.89 ± 0.01 g lactate/g glucose (∼90% of the maximum theoretical yield). In contrast, when growing on xylose, B-768 exhibited a much slower growth rate of 0.32 ± 0.01 h^-1^. After ∼24 h, cells reached the growth of 3.45 ± 0.10 OD_600_ and only consumed 3.52 ± 0.45 g/L out of ∼10 g/L xylose available with a xylose uptake rate of 0.35 ± 0.04 g xylose/OD_600_/h (Fig. 5A, 5B). B-768 only produced lactate at a titer of 1.57 ± 0.12 g lactate/L, a productivity of 0.07 ± 0.01 g lactate/OD_600_/h, and a yield of 0.18 ± 0.02 g lactate/g xylose. Overall, faster growth and glucose uptake rate made B-768 reroute high carbon flux toward lactate biosynthesis. Even though B-768 reached the same cell biomass for growth on xylose after 24 h, slower growth and xylose uptake rate shifted its carbon flux mainly towards biomass synthesis at the expense of lactate production.

**Figure 5:**
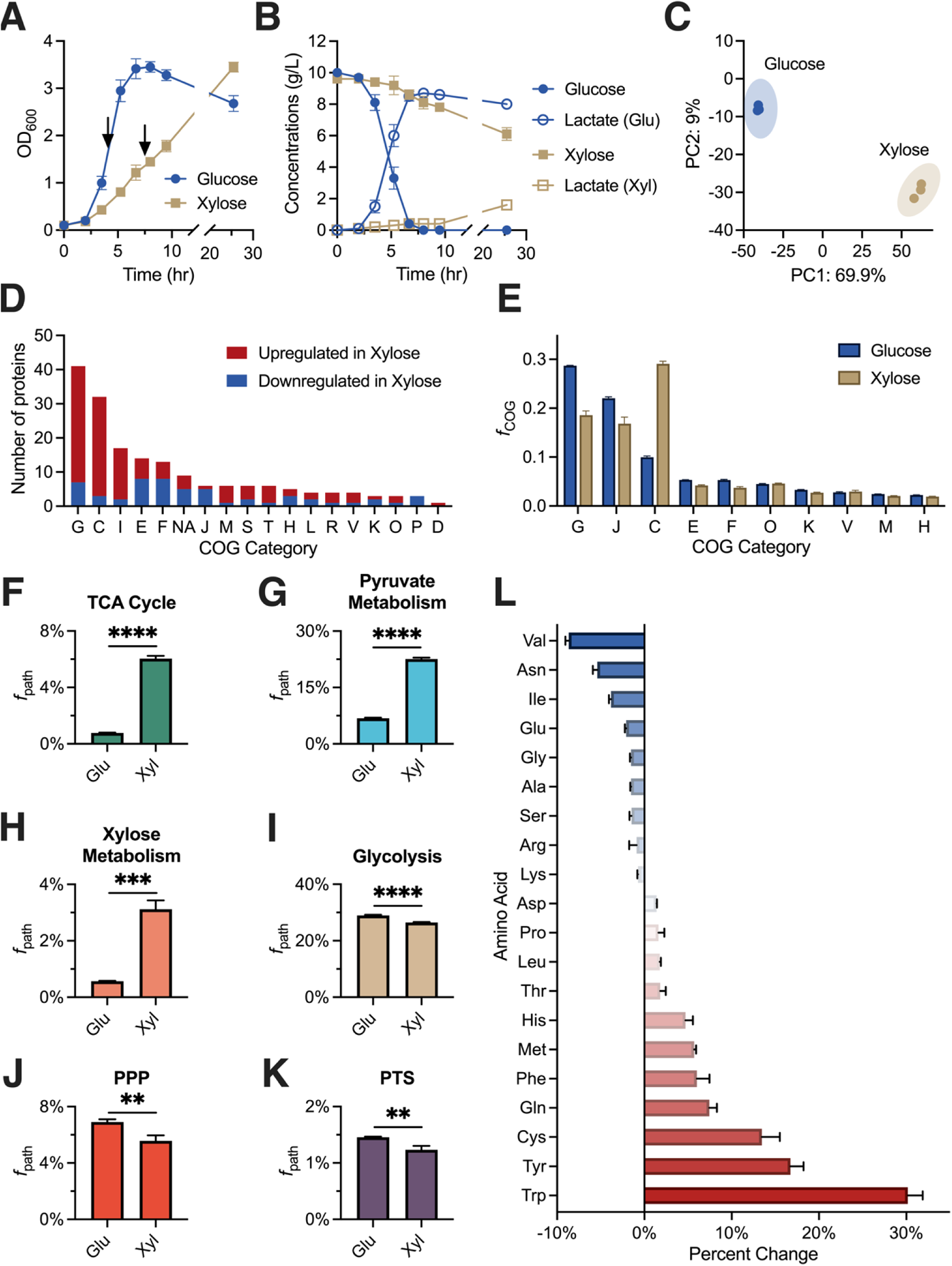
*B. coagulans* B-768 exhibited distinct proteomes for growing on glucose and xylose. (A–B) Kinetic profiles of (A) cell growth and (B) glucose/xylose consumption and lactate production. Proteomics samples were collected from parallel cultures at the time points indicated by black arrows. (C) Principal component analysis of *B. coagulans* B-768 proteomes for growth on glucose and xylose. (D) Number of differentially expressed proteins between glucose and xylose conditions for most perturbed COG categories. (E) Mass fraction of proteomes in top COG categories (*f*_COG_) for glucose and xylose conditions. (F–K) Proteome allocation by pathway for glucose and xylose conditions. (L) Percent change in amino acid allocation of proteomes between glucose and xylose conditions. COG Categories: G = Carbohydrate Transport and Metabolism; C = Energy Production and Conversion; I = Lipid Transport and Metabolism; E = Amino Acid Transport and Metabolism; F = Nucleotide Transport and Metabolism; NA = No COG annotation; J = Translation; M = Cell Wall/Membrane/Envelope Biogenesis; S = Function Unknown; T = Signal Transduction Mechanisms; H = Coenzyme Transport and Metabolism; L = Replication, Recombination, and Repair; R = General Function Prediction Only; V = Defense Mechanisms; K = Transcription; O = Posttranslation Modification, Protein Turnover, Chaperones; P = Inorganic Ion Transport and Metabolism; D = Cell Cycle Control, Cell Division, Chromosome Partitioning. Statistics for F–K were performed using an unpaired t-test with default settings in GraphPad Prism v10.2.3. Error bars represent the standard deviation of 3 biological replicates. **** = p-value < 0.0001; *** = p-value < 0.001; ** = p-value < 0.01.

#### B-768 exhibited distinct proteomes for growth on glucose and xylose

To elucidate the phenotypic differences for B-768 grown on glucose and xylose at the cellular level, cultures were sampled at mid-exponential phase for quantitative proteomics. Principal component analysis of glucose and xylose samples revealed highly distinct proteomes (Fig. 5C). Differential expression analysis was then performed on the glucose-and xylose-only samples and the number of significantly altered proteins was tallied for each COG category (Fig. 5D). The results showed that proteomes had shifted across the COG categories to varying degrees. Most significantly perturbed proteins in cells growing on xylose compared to on glucose belong to the Carbohydrate Transport and Metabolism (COG-G) and Energy Production and Conversion (COG-C) categories. As expected, both the major xylose utilization proteins—xylose isomerase and xylulokinase—were among the most xylose-upregulated proteins by 5.99 log_2_-fold (q-value = 4.17E-14) and 7.28 log_2_-fold (q-value = 4.17E-14), respectively in the COG-G category (Fig. 4, Dataset 3). Among the PPP proteins, ribose-5-phosphate isomerase (RPI) was significantly upregulated by 1.29 log_2_-fold (q-value = 0.0478), while the rest were not significantly perturbed (Fig. 4, Dataset 3). Most of the xylose downregulated proteins in COG-G were proteins involved in the EMP pathway, such as triose phosphate isomerase (TPI), glyceraldehyde-3-phosphate dehydrogenase (GAPDH), enolase (ENO), and pyruvate kinase (PK) by -1.9 log_2_-fold (q-value = 8.45E-3), -1.29 log_2_-fold (q-value =3.53E-3), -1.36 log_2_-fold (q-value = 0.012), and -1.48 log_2_-fold (q-value = 2.93E-4), respectively (Fig. 4, Dataset 3). Lactate dehydrogenase (LDH) is also significantly downregulated in xylose relative to glucose by -2.63 log_2_-fold (q-value = 1.83E-4) at the measured timepoint (Fig. 4, Dataset 3), which is consistent with the observed low lactate production (Fig. 5B). In COG-C, several enzymes involved in pyruvate metabolism, such as formate C-acetyltransferase (or pyruvate formate lyase, PFL), dihydrolipoamide dehydrogenase (DLD), and pyruvate dehydrogenase subunits E1⍺ and E1β (PDH) for converting pyruvate to acetyl-CoA, were significantly upregulated in xylose by 5.71 log_2_-fold (q-value = 4.17E-14), 3.71 log_2_-fold (q-value = 4.17E-14), 3.44 log_2_-fold (q-value = 2.52E-6), and 3.1 log_2_-fold (q-value = 1.37E-13), respectively (Dataset 3). Enzymes that convert acetyl-CoA into acetate and ethanol were upregulated in xylose, including acetyl-CoA synthetase (ACS), aldehyde dehydrogenase (ALDH), and alcohol dehydrogenase (ADH), by 3.7 log_2_-fold (q-value = 1.24E-4), 4.86 log_2_-fold (q-value = 4.17E-14), and 3.6 log_2_-fold (q-value = 1.03E-7), respectively (Dataset 3 and Fig. S4). Acetyl-CoA is also passed into the TCA cycle, which is highly upregulated on xylose (Dataset 3 and Fig. S5).

#### Tradeoffs of proteome reallocation between carbohydrate metabolism and energy generation contributed to the differential growth on glucose and xylose

We then investigated how B-768 re-allocated its resources in response to glucose or xylose by summing the mass fractions of all expressed proteins for each COG (Fig. 5E). The top 7 COGs, including COG-G, J (Translation), C, E (Amino Acid Transport and Metabolism), F (Nucleotide Transport and Metabolism), O (Posttranslation Modification, Protein Turnover, Chaperons), K, V (Defense Mechanisms), M (Cell Wall/Membrane/Envelope Biogenesis), and H (Coenzyme Transport and Metabolism), contributed up to 80% of the total measured proteomes as shown in Fig. 5E. The top 3 COGs, including G, J, and C, shared up to 60–65% of each measured proteome with a high degree of variability for growth on glucose and xylose. Despite most significantly differential proteins in COG-G being upregulated in xylose relative to glucose (Fig. 5D), the proteome mass fraction allocated to COG-G was much higher when B-768 was grown on glucose compared to xylose (29% Glu vs. 19% Xyl, Fig. 5E). This result indicates that protein expression was higher even with fewer proteins in cells grown in glucose compared to cells grown in xylose. Consistent with this hypothesis, B-768 also devoted significantly more resources to COG-J when grown on glucose compared to xylose (22% Glu vs. 17% Xyl). Xylose-grown cells disproportionately invested their resources in COG-C, presumably to offset the higher energy cost of transporting and assimilating xylose (29% Xyl vs. 10% Glu, Fig. 5E).

To better elucidate how proteome reallocation is linked to the distinct phenotypes for B-768 growing on glucose and xylose, the allocation of proteomes in COG-G and COG-C was analyzed at a more granular resolution based on the relevant pathways (Fig. 5F–K). Enriched pathways in the cells grown on xylose include the TCA cycle (6.78-fold, p-value < 0.0001, Fig. 5F), pyruvate metabolism (2.32-fold, p-value < 0.0001, Fig. 5G), and xylose metabolism (4.52-fold, p-value = 0.0001, Fig. 5H), reiterating the trends observed from differential expression analyses. Glucose-grown cells had slightly more investments into glycolysis (0.095-fold, p-value < 0.0001, Fig. 5I), the pentose phosphate pathway (0.24-fold, p-value = 0.0063, Fig. 5J), and the phosphotransferase system (0.18-fold, p-value = 0.0054, Fig. 5K). Overall, these findings reveal that B-768 redistributed its proteome to accommodate the increased energy demand of metabolizing xylose. To do this, various carbohydrate metabolism and energy production genes were overexpressed, which direct carbon flux from xylose through the PPP to yield pyruvate. Pyruvate is then converted into energy by channeling carbon and electron fluxes through pyruvate metabolism, then the TCA cycle, followed by the electron transport system for ATP synthesis. Although ATP synthase was not significantly different between xylose and glucose conditions, other key proteins in the electron transport system were upregulated in xylose, such as NADH dehydrogenase by 8.93 log_2_-fold (q-value = 3.63E-9) and succinate dehydrogenase by 3.72 log_2_-fold (q-value = 2.42E-5). A significant reduction in LDH expression and lactate production, and an increase in pyruvate metabolism correlates with the decreased lactate metabolism for growth on xylose.

Lastly, amino acid reallocation in the measured proteomes was performed as previously described to investigate the metabolic cost associated with switching to a xylose-utilizing proteome(64) (Fig. 5L). Higher amounts of costly amino acids like tryptophan, tyrosine, and phenylalanine are used in xylose-only conditions compared to glucose-only conditions(65). Consistent with this finding, the 10 most highly expressed proteins in xylose possess 107% more tryptophan, 37% more tyrosine, and 21% more phenylalanine than the top 10 proteins for glucose. The greater demand for these costly amino acids could help explain the marked investment into energy generation processes when B-768 is grown on. This finding suggests that supplementation of *B. coagulans* B-768 with metabolically expensive amino acid(s) may alleviate part of the metabolic burden due to amino acid biosynthesis and improve growth on xylose.

### Carbon catabolite repression is dominant in B-768 growing on glucose-xylose mixture and switchgrass biomass hydrolysate

As cellulosic hydrolysates contain a mixture of C5 and C6 sugars, it is important to understand how B-768 regulates its metabolism for growth on mixed sugars. Carbon catabolite repression (CCR) is a common regulatory mechanism used by bacteria to inhibit metabolism of sub-optimal carbon sources (e.g., xylose) in the presence of a preferred substrate (e.g., glucose). The diauxic growth observed when B-768 is grown on either a mixture of refined glucose and xylose (G+X) or switchgrass hydrolysate (SGH), is evidence for a strong CCR on xylose assimilation by glucose (Figs. 6A, 6B). Consistent with this observation (Fig. 6B), expression of the *xyl* operon was heavily repressed (Figs. 6D, 6F). The effects of the observed mechanism can be captured at the cellular level. Principal component analysis shows that all proteomes of cultures growing in the presence of glucose were clustered tightly together, while those grown on xylose were highly diverged (Fig. 6C).

**Figure 6:**
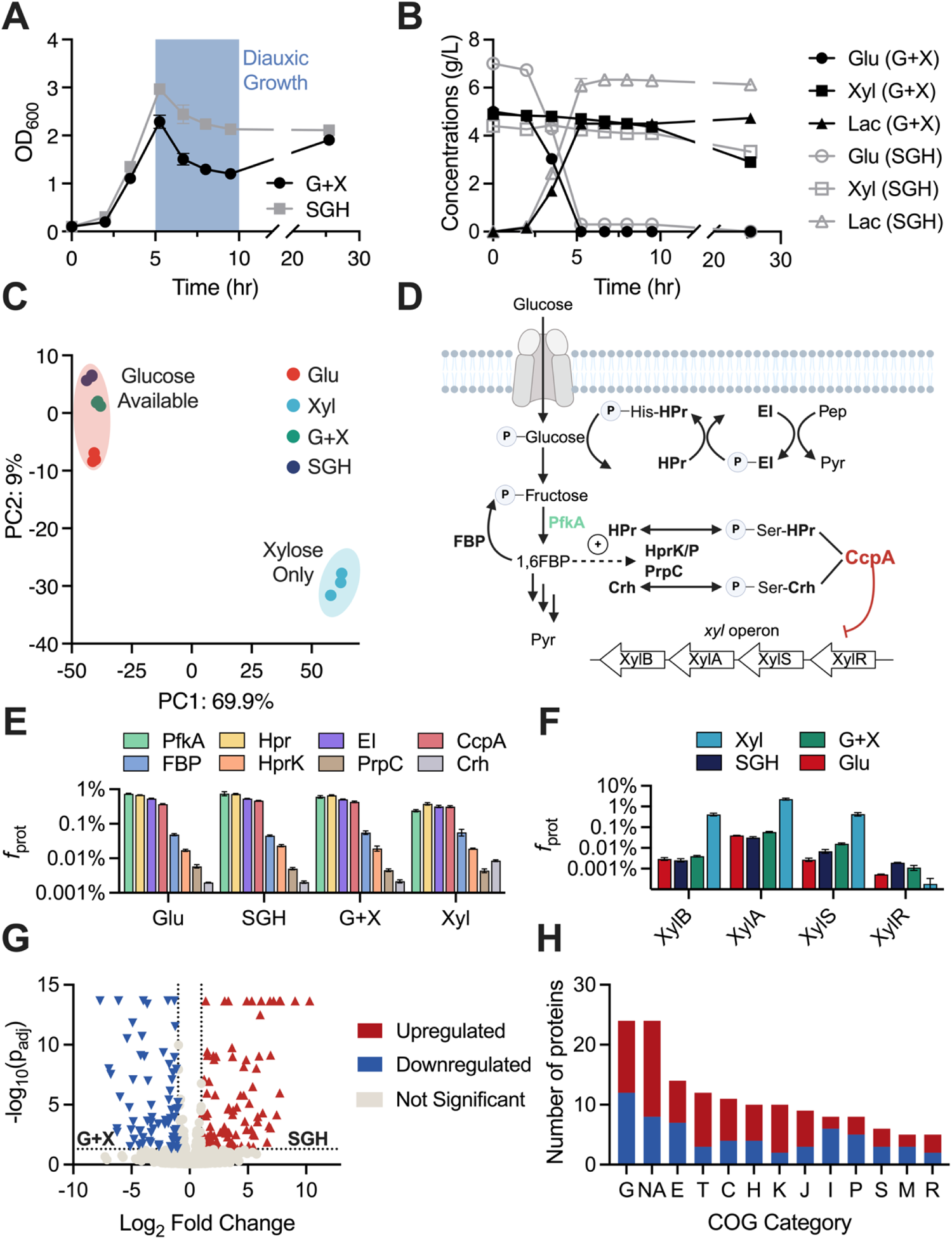
Growth of B-768 in refined sugars and raw hydrolysate reveals mechanisms of cellular robustness and carbon catabolite repression. (A) Growth profiles of B-768 grown on glucose + xylose (G+X) mixture and switchgrass hydrolysate (SGH). (B) Glucose and xylose consumption and lactate production profiles of B-768 grown on G+X and SGH. (C) Principal component analysis plot of B-768 proteomes grown on different sugars. (D) Schematic of CCR in *Bacillus* species. (E) Levels of CCR proteins in B-768 proteomes grown on different sugars. (F) Levels of *xyl* proteins in B-768 proteomes grown on different sugars. (G) Volcano plot of differentially expressed proteins between SGH and G+X conditions. (H) Number of significantly perturbed proteins by COG category. COG Categories: G = Carbohydrate Transport and Metabolism; E = Amino Acid Transport and Metabolism; T = Signal Transduction Mechanisms; C = Energy Production and Conversion; H = Coenzyme Transport and Metabolism; NA = No COG annotation; K = Transcription; J = Translation; I = Lipid Transport and Metabolism; P = Inorganic Ion Transport and Metabolism; S = Function Unknown; M = Cell Wall/Membrane/Envelope Biogenesis; R = General Function Prediction Only.

To characterize CCR in *B. coagulans* B-768 and to select genetic targets for eliminating it, canonical CCR proteins were quantified across a variety of growth conditions based on the CCR model of the well-studied *B. subtilis* (Figs. 6D, 6E). To enable CCR, the master regulator CcpA and its primary corepressor, histidine-containing phosphocarrier protein (HPr), or the secondary catabolite corepressor (Crh), bind to catabolite-responsive elements (*cre*) located in the open reading frame or promoter region of the target genes and block transcription(66) (Fig. 6D). Hpr and Crh bind CcpA when phosphorylated at the Ser46 residue by the bifunctional HPr kinase/phosphorylase (HPrK/P), whose activity is increased by the glycolytic intermediate, fructose-1,6-bisphosphate (FBP)(67–69). CCR is further modulated by glucose-6-phosphate (G6P) and FBP, which improve the DNA binding affinity of CcpA-(Hpr-Ser46-P), but not CcpA-(Crh-Ser46-P)(70, 71). The results show that protein levels of CcpA, HPr, and HprK/P did not significantly change between glucose-available and xylose-only conditions, agreeing with a previous report (Fig. 6E)(72). While not statistically significant, phosphofructokinase-1 (PfkA), which converts fructose-6-phosphate into FBP, is downregulated by 1.47 log_2_-fold changes in xylose cultures relative to glucose. For growth on xylose, the FBP node of the EMP pathway can be effectively bypassed because carbon passed through the PPP connects to glycolysis downstream of FBP at glyceraldehyde-3-P. Lower concentrations of FBP would in turn downregulate the kinase activity of HprK and DNA binding affinity of the CcpA effector complex (Fig. 6D). Therefore, in B-768 relief of CCR on xylose appears to be likely due to reduced intracellular levels of FBP.

### Growth hardiness of *B. coagulans* B-768 on complex biomass hydrolysate is explained by proteomics

A major challenge to utilizing lignocellulosic sugars is overcoming microbially inhibitory chemicals formed during pretreatment and hydrolysis of the biomass. Depending on their concentration, they can lead to extended lag phases, reduced product yields, or stalled fermentations. While there are methods to mitigate these inhibitors, their drawbacks included added cost, sugar loss, and reduced overall conversion efficiency(73, 74). Therefore, using microbial biocatalysts that are robust to the presence of inhibitory compounds is paramount(75). To elucidate the cellular robustness of B-768, its growth and lactate production were compared for undetoxified dilute-acid switchgrass hydrolysate and a refined sugar control (Figs. 6A, 6B). Surprisingly, no inhibitory effect on growth and lactate production was detected for B-768 grown on SGH compared to refined sugars. This result indicates that B-768 can utilize glucose and—to a much lesser extent—xylose present in undetoxified biomass hydrolysate without a reduction in growth or lactate production, which makes this strain particularly attractive as a microbial biomanufacturing platform.

To investigate the molecular drivers for the observed cellular robustness, comparative proteomics was performed between cells grown in SGH and X+G (Fig. 6G). COG analysis was performed on the 161 significantly differential proteins and all but 24 were assigned to a COG category (Fig. 6H). Most proteins that could not be assigned a COG were annotated as hypothetical proteins, but a few notable proteins included a putative transpose, a putative type VII secretion system protein, a flagellar protein, and two sporulation proteins. All these proteins are stress-implicated and probably play a role in helping B-768 adapt to hydrolysate toxicity. The top 3 COG categories were COG-G (n = 24), COG-E (n = 14), and COG-T (n = 12). In terms of carbon metabolism, all 3 major arabinose utilization proteins, L-arabinose isomerase, L-ribulokinase, and L-ribulose-5-P 4-epimerase were significantly upregulated in SGH relative to X+G by 5.33 log_2_-fold (q-value = 2.15E-14), 6.82 log_2_-fold (q-value = 5.98E-4), and 7.47 log_2_-fold (q-value =6.88E-5), respectively. Additionally, 6-phospho-β-glucosidase (cellobiose utilization) and galactokinase (galactose utilization) were upregulated in SGH by 9.06 log_2_-fold (q-value = 2.15E-14) and 4.07 log_2_-fold (q-value = 1.17E-4), respectively (Dataset 4). This result indicates that B-768 actively metabolizes arabinose, cellobiose, and galactose when grown on unrefined biomass hydrolysates. In addition to these proteins, several transporters of both the ABC and PTS varieties (n = 9), and chemotaxis-implicated proteins (n = 4) were significantly altered (Table 5). Similar proteins were previously found to be significantly perturbed in *Clostridium acetobutylicum* in response to phenolic inhibitors—common toxins found in biomass hydrolysates(76, 77). Another inhibitor found in biomass hydrolysate is methylglyoxal(78, 79); therefore, it is noteworthy that the most upregulated protein in SGH relative to X+G was glyoxylase II, a crucial enzyme for detoxifying methylglyoxal(80). Finally, 10 putative transcriptional regulators were differentially expressed (8 up, 2 down) with actuate transcriptional changes to accommodate growth in hydrolysates (Table 5). Together, these results provide promising engineering targets for further improving B-768’s exceptional hydrolysate tolerance and demonstrate that B-768 displays a robust and diverse proteomic response to hydrolysate toxicity, enabling favorable growth kinetics and lactate titers in comparison to other industrial biocatalysts.

**Table 5:** Differential protein expression for candidate proteins contributing to cellular robustness of B-768 growing in SGH.

| Protein ID | Function | Class | Log<sub>2</sub>-fold Change | q-value |
| --- | --- | --- | --- | --- |
| Ga0586967_01_1426112_1427749 | methyl-accepting chemotaxis protein | Chemotaxis | 7.21 | 2.2E-14 |
| Ga0586967_01_2953390_2954649 | methyl-accepting chemotaxis protein | Chemotaxis | 2.1 | 0.00377 |
| Ga0586967_01_1922219_1923952 | methyl-accepting chemotaxis protein | Chemotaxis | 2.02 | 0.0171 |
| Ga0586967_01_3602104_3603432 | methyl-accepting chemotaxis protein | Chemotaxis | -6.24 | 0.00197 |
| Ga0586967_01_1885126_1886544 | PTS system trehalose-specific IIC component | Transport | -1.06 | 0.0103 |
| Ga0586967_01_1886575_1887057 | PTS system glucose-specific IIA component | Transport | -1.18 | 0.00121 |
| Ga0586967_01_1467232_1467561 | PTS system cellobiose-specific IIA component | Transport | -1.38 | 0.00901 |
| Ga0586967_01_1541669_1542151 | fructoselysine and glucoselysine-specific PTS system IIB component | Transport | -2.64 | 0.0316 |
| Ga0586967_01_2529421_2529912 | mannose/fructose/N-acetylglactosamine-specific phosphotransferase system component IIB | Transport | -3.63 | 0.00023 |
| Ga0586967_01_1189660_1191024 | PTS system ascorbate-specific IIC component | Transport | -3.89 | 9.7E-10 |
| Ga0586967_01_1452905_1453891 | LacI family transcriptional regulator | Regulator | 7.71 | 2.2E-14 |
| Ga0586967_01_1692988_1693203 | putative transcriptional regulator | Regulator | 5.56 | 0.00345 |
| Ga0586967_01_633467_634027 | TetR/AcrR family transcriptional repressor of lmrAB and yxaGH operons | Regulator | 3.97 | 0.00309 |
| Ga0586967_01_3908323_3909216 | phenylacetic acid degradation operon negative regulatory protein | Regulator | 3.62 | 0.0132 |
| Ga0586967_01_2622621_2623172 | DNA-binding transcriptional ArsR family regulator | Regulator | 3.09 | 0.00232 |
| Ga0586967_01_1173785_1174315 | DNA-binding MarR family transcriptional regulator | Regulator | 2.63 | 0.0019 |
| Ga0586967_01_30811<br>24_3082011 | DNA-binding transcriptional<br>LysR family regulator | Regulator | 2.27 | 9.3E-10 |
| Ga0586967_01_33636<br>33_3364772 | GntR family transcriptional<br>regulator of arabinose operon | Regulator | 2.03 | 7.9E-10 |
| Ga0586967_01_19153<br>88_1915990 | AcrR family transcriptional<br>regulator | Regulator | -3.24 | 0.00012 |
| Ga0586967_01_35848<br>55_3586105 | DNA-binding transcriptional<br>ArsR family regulator | Regulator | -4.86 | 8.4E-08 |

## CONCLUSIONS

Through detailed phenotype characterization, comparative functional genomics, and quantitative proteomic analyses, this study elucidates the mechanisms underlining the robustness of undomesticated *B. coagulans* strain B-768,. Highlighted is the metabolic capability of B-768 to utilize a wide range of C5 and C6 sugars at various temperatures, with glucose as the preferential substrate. We created a high-quality genome assembly and annotation of *B. coagulans* B-768. Essential to the robustness of B-768 is its larger genome including the expanded Carbohydrate Transport and Metabolism and Mobilome, which enables improved sugar utilization and horizontal gene transfer. A large set of genes devoted to its innate and adaptive defense mechanisms should help *B. coagulans* resist phage attacks, which are a risk for bacterial industrial fermentation processes. B-768 exhibits distinct proteomes for growth on glucose and xylose, consistent with its observed phenotypes. Faster growth and glucose uptake rate resulted in lactate overflow metabolism while slower growth and xylose uptake rate diminished lactate production. This difference could be explained by the tradeoff in proteome reallocation between the Carbohydrate Transport and Metabolism process and the Energy Generation and Production process. B-768 can utilize biomass hydrolysates with various C5 and C6 sugars controlled by a tightly regulated gene network. Overall, B-768 is a robust microbial biomanufacturing platform that has potential beyond lactate fermentation. A fundamental understanding of its robustness revealed in this study will be critical for rechanneling and tuning its metabolism for optimized conversion of lignocellulosic biomass to chemicals, fuels, and materials beyond lactate.

## MATERIALS AND METHODS

### Cell Culture

#### Strains

*B. coagulans* strain B-768 (ATCC 10545=NRS-784) was obtained from the NRRL Agriculture Research Service strain collection of the United States Department of Agriculture (USDA) located in Peoria, IL. The strain was stored as glycerol stocks at -80 °C.

#### Media

The basal medium contained 1.5 g/L yeast extract, 100 mM sodium MES buffer (pH 6.5), and mineral solution. The mineral solution contained: 2 g/L (NH_4_)_2_SO_4_, 2 g/L KH_2_PO_4_, 2 g/L NaCl, 0.2 g/L MgSO_4_·7H_2_O, 5 mg/L MnSO_4_·H_2_O, and 10 mg/L FeSO_4_·7H_2_O. The phosphate buffered saline (PBS) contained per liter: 8 g NaCl, 0.2 g KCl, and 1.44 g Na_2_HPO_4,_ adjusted to pH 7. 4 with NaOH.

The switchgrass hydrolysate (SGH) medium contained per liter: 10% (v/v) of concentrated dilute acid pretreated SGH and 100 mM MES buffer (pH 6.5). The concentrated SGH solution was prepared as previously described(81), which contained: ∼70 g/L glucose and ∼44 g/L xylose as the main C5/C6 sugars of biomass hydrolysates. SGH was used at 10%w/w for bottle cultures.

#### High-throughput Microplate Growth Kinetic Experiments

Growth of B-768 on various carbon sources (glucose, xylose, cellobiose, galactose, mannose, or arabinose) at various temperatures (37, 40, 45, 47, 53, or 55 °C) was measured high-throughput using Bioscreen Pro™ (Oy Growth Curves Ab Ltd., Finland). B-768 was inoculated in basal medium containing no sugar and grown at 50 °C overnight. Next day, overnight grown cells were diluted in fresh medium with initial OD_600_ of 0.1 in total 300 µL and growth was monitored at 580 nm for 24 or 48 hrs. For the lactate tolerance experiment, cells were prepared in a basal medium with 8 g/L glucose, and growth was monitored at 50 °C using the Bioscreen Pro™.

#### Bottle Cultures

Seed cultures and media were prepared as described for the high-throughput studies. Fermentations were conducted in 25 mL Pyrex bottles (Corning Life Sciences, NY) filled with 20 mL of medium and incubated and mixed at 50 °C and 100 rpm, respectively. Cultures were sampled for sugars (glucose and xylose), lactate, and OD_600_ using a 1 mL sample. Proteomic samples (1 mL) were withdrawn, centrifuged, washed with chilled dH_2_O, and the pellet was stored at -80 °C.

#### pH-controlled Fermentations

Fermentations were maintained at pH 6.5 and 50 °C (unless stated otherwise) and stirred at 300 rpm. Cells were inoculated to an initial OD_600_ of 0.1. The pH was maintained by adding 4 N KOH. Cultures were sampled for OD_600_, sugars, and fermentation products.

### Analytical Methods

#### High-Performance Liquid Chromatography (HPLC)

Metabolites such as sugars and acids (e.g. lactate and acetate) were measured by Bio-Rad HPX-87H column at 65 °C with 5 mM sulfuric acid as a mobile phase using a Thermo high-performance liquid chromatography (Ultimate 3000) system equipped with a refractive index (RI).

#### Liquid Chromatography with Tandem Mass Spectrometry (LC-MS/MS)

*B. coagulans* strain B-768 cultured in the presence of various sugars (n = 3 per glucose, xylose, a mixture of glucose and xylose, and switchgrass hydrolysate) were collected by centrifugation and cell pellets prepared for LC-MS/MS based proteomics as previously described(82). All raw mass spectra for quantification of proteins used in this study have been deposited in the MassIVE and ProteomeXchange data repositories under accession numbers MSV000095240 (MassIVE) and PXD053622 (ProteomeXchange), with data files available at ftp://massive.ucsd.edu/v08/MSV000095240/.

#### Genome Sequencing

Genomic DNA of B. coagulans B-768 was subjected to Pacific Biosciences (PacBio) long-read sequencing by DOE Joint Genome Institute (JGI)(83). A >10 kb Pacbio SMRTbell^®^ library was constructed and sequenced on the PacBio Sequel platform, which generated 78,854 high-fidelity CCS reads totaling 696,902,661 bp. Reads >5kb were assembled with Flye v2.8.3 using default settings(84, 85). The final draft assembly contained 1 contig in 1 scaffold, totaling 3,944,476 bp in size, and was 99.95% complete according to CheckM(86). The input read coverage was 181.4x. The final draft assembly was annotated using the Integrated Microbial Genomes Annotation Pipeline (IMGAP) v5.1.1. The whole-genome assemblies and annotation were deposited at DDBJ/EMBL/GenBank (Bioproject ID: PRJNA1123721).

### Bioinformatics and Data Analysis

#### CRISPR System and Spacer Identification

CRISPR system prediction was performed using CRISPRCasTyper v1.8.0 with default settings(87). For identification of spacer hits on plasmids, nucleotide sequences for the entire NCBI collection of bacterial plasmid sequences were downloaded from PLSDB v. 2021_06_23_v2(88) and used to construct a BLASTn database against which the JGI-predicted CRISPR spacers were blasted using the blastn-short algorithm with default settings. Hits with e-value < 0.05 were retained and the top hit for each spacer was chosen as the target source. For identification of spacer hits on phages, SpacePHARER v5.c2e680a(89) was used with the GenBank_phage_2018_09 database and default settings.

#### Restriction-Modification Annotation

All B-768 protein sequences were searched against the complete REBASE database using BLASTp v2.15.0+ with default settings. Hits were filtered down to those with a percent identity >60% and alignment coverage >80% (alignment length/query length). Hits were then sorted by ascending e-value and the top hit for each query was retained. 11 hits remained and all had e-values < 7.5E-42. These were considered bonafide R-M homologs and the REBASE classification was used to assign the type.

#### Circular Genome Map

Proksee v5.2.1(90) was used to generate a circular genome map for B. coagulans B-768 from functional annotation provided by JGI. GC content and skew were calculated using the built-in function in Proksee. COG mapping was performed manually using the feature upload tool and the output from COGclassifier v1.0.5(91). Prophage regions were predicted using PHASTEST v3.0 and mapped using the feature upload tool(40).

#### Pan-genome Phylogeny

Genome sequences of all RefSeq B. coagulans strains, except RUG1433, were downloaded from NCBI. A core-genome alignment was performed on the pan-genome using Parsnp v1.7.4 with default settings(92). A recombination-aware maximum likelihood tree was then constructed from the core-genome alignment using Gubbins v3.2.1 with default settings(93).

#### Proteomics

The resulting MS/MS spectra acquired from the Q Exactive MS were searched against the B. coagulans B-768 proteome database appended with common protein contaminants using the SEQUEST HT algorithm in Proteome Discoverer v.2.5 (Thermo Scientific) and peptide spectrum matches (PSMs) scored and filtered with Percolator (settings and dynamic/static mods described in Walker et al’s study(82)). Peptides (FDR < 1%) were then quantified by chromatographic peak area and derived abundances mapped back to each peptide’s respective protein and peptide abundance summed to estimate protein-level abundances. Protein abundances were then log_2_ transformed, normalized by LOESS, and median centered using InfernoRDN(94). Missing values were imputed, and statistical tests (T-test and ANOVA) performed using Perseus v2.0.7.0(95). Log_2_ transformed and imputed protein abundance data were analyzed using the DEP R package v1.24.0(96). Significant proteins were filtered out based on a 5% FDR and minimum log_2_ fold-change of 1.

#### Proteome Reallocation

Based on the mass fractions of each protein per sample, proteome reallocation was calculated by dividing each raw protein abundance by the sum of all protein abundances for a given sample and summed up across the indicated categories of interest(64, 82). COG annotations were mapped onto predicted protein sequences from JGI using COGclassifier v1.0.5(91). All calculations were performed in R v4.3.0.

#### Amino Acid Allocation

Analyses of amino acid allocation in the measured proteomes were determined based on the formulation previously described(64, 82). All the calculations were performed in Python v3.8.16 and the outputs were analyzed in R v4.3.0.

#### Kinetics

The specific growth rate was determined by assuming that cell growth follows the first-order kinetics (Equation 1).

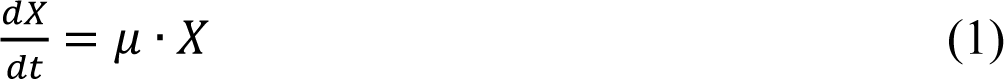

where X (g DCW/L) is cell mass, t (h) is the culturing time, and μ (1/h) is the specific growth rate. Note that DCW represents dry cell weight and is linearly correlated with the measured cell optical density (OD_600_). The substrate uptake rate was determined as follows (Equation 2):

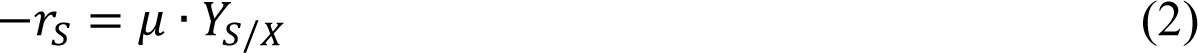

where r_S_ (g/g DCW/h) is the specific substrate uptake rate and Y_S/X_ (g substrate/g CDW) is the yield of a substrate (e.g., glucose) to cell mass. The product formation rate was determined as follows (Equation 3):

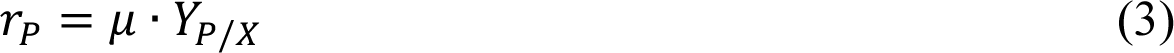

where r_P_ (g/g DCW/h) is the specific substrate uptake rate and Y_P/X_ (g substrate/g CDW) is the yield of a product (e.g., lactate) to cell mass.

#### Calculation of Optimal Growth Temperatures

Optimal growth temperatures on various sugars were determined by fitting maximum growth rate versus temperature with a non-linear model (Continuous hinge function. Segmental regression lines with gentle connection, least squares fit) using GraphPad Prism v10.2.3.

## ACKNOWLEDGEMENTS

This research was funded by the DOE BER award (DE-SC0022226 to CTT, RJG, BD). The sequencing project was supported by the Center for Bioenergy Innovation (CBI), U.S. Department of Energy Bioenergy Research Centers, funded by the Office of Biological and Environmental Research in the DOE Office of Science. The work (Project IDs # 510751, 509417) conducted by the U.S. Department of Energy Joint Genome Institute, a DOE Office of Science User Facility, is supported under Contract No. DE-AC02-05CH11231. Dien, Jackson, and Slininger received financial support from the U.S. Department of Agriculture, Agricultural Research Service, United States (CRIS Numbers 5010-41000-189). The mention of trade names or commercial products in this article is solely for the purpose of providing scientific information and does not imply recommendation or endorsement by the U.S. Department of Agriculture. USDA is an equal opportunity provider and employer.

