## Supplementary Materials for "Expanded Genome and Proteome Reallocation in a Novel, Robust *Bacillus coagulans* Capable of Utilizing Pentose and Hexose Sugars"

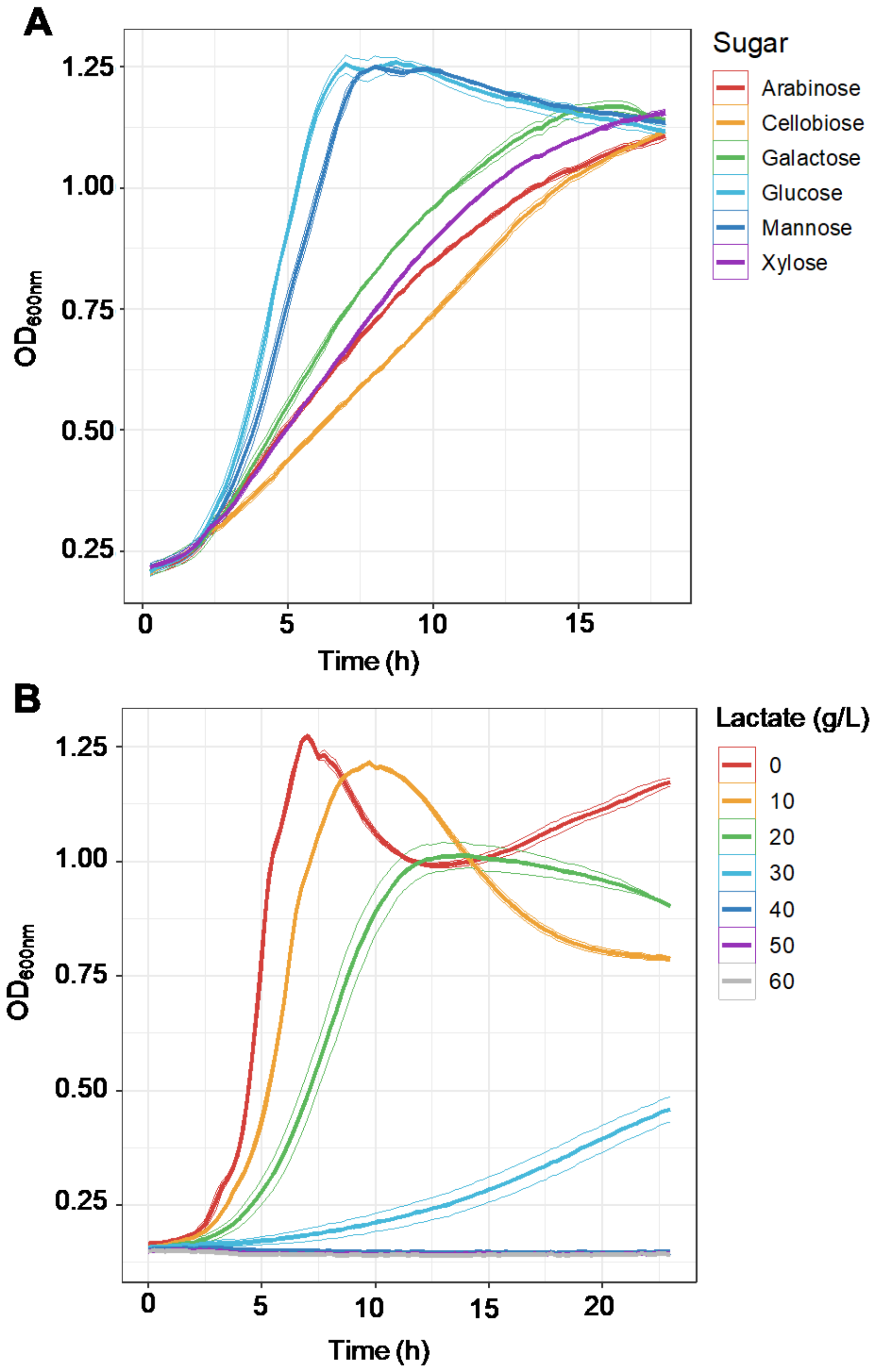


**Figure S1:** Growth kinetics of *B. coagulans* B-768 growing **(A)** on various C5 and C6 sugars and **(B)** in presence of various concentrations of lactate at 50°C.

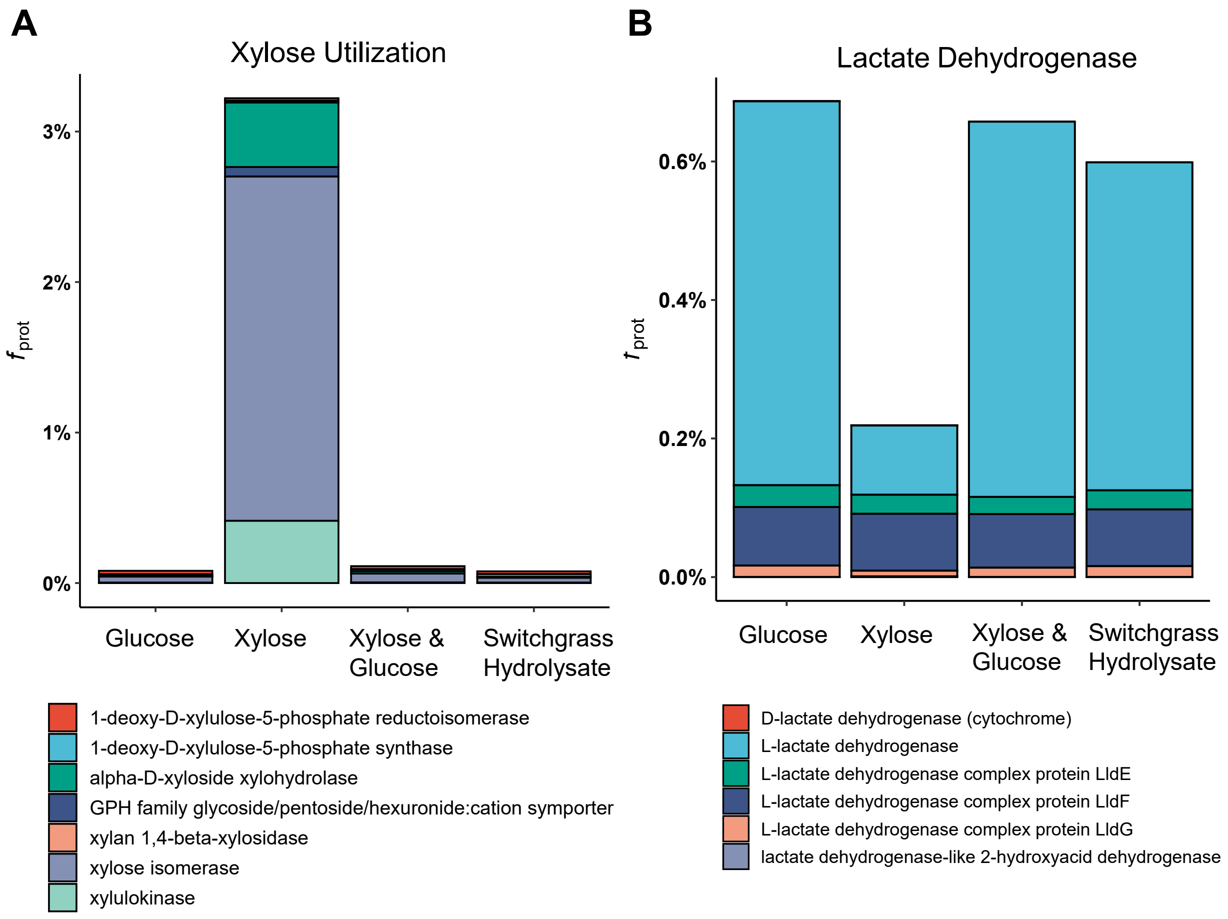


**Figure S2:** Proteome reallocation for **(A)** xylose utilization and **(B)** lactate biosynthesis of *B. coagulans* B-768 growing xylose, glucose, a mixture of glucose and xylose, and switchgrass hydrolysate (SGH).

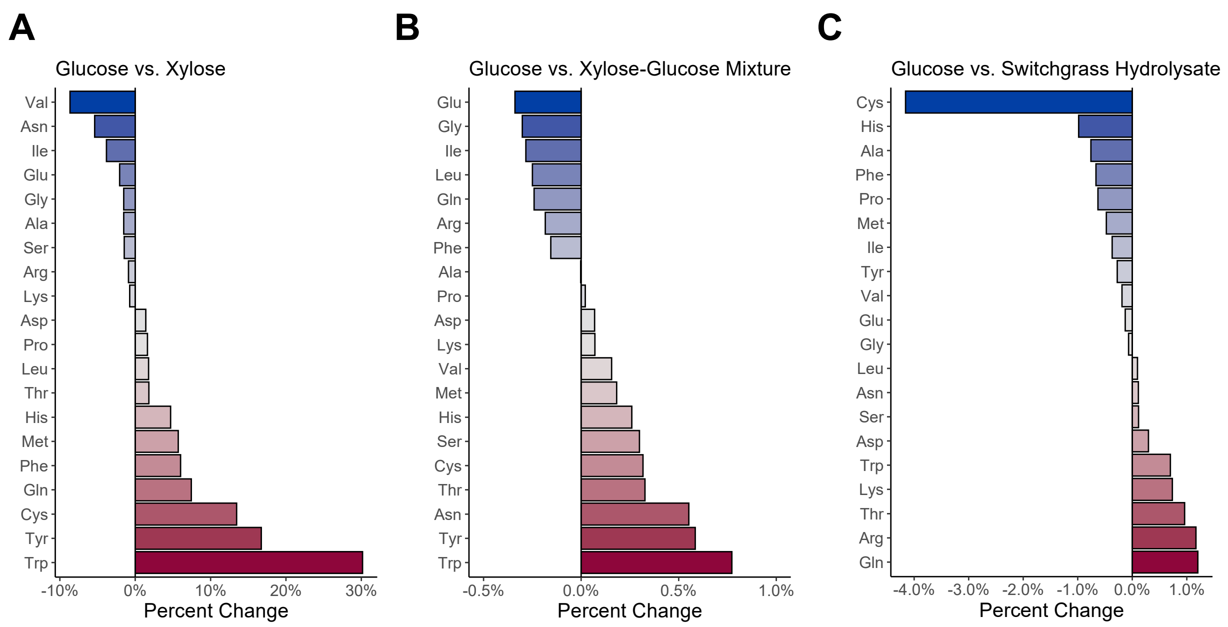


**Figure S3:** Percent of change in amino acid reallocation in measured proteomes of *B. coagulans* B-768 growing on various sugars including **(A)** glucose versus xylose, **(B)** glucose versus a mixture of glucose and xylose, and **(C)** glucose versus switchgrass hydrolysate (SGH).

### Differential Expression of Xylose vs. Glucose

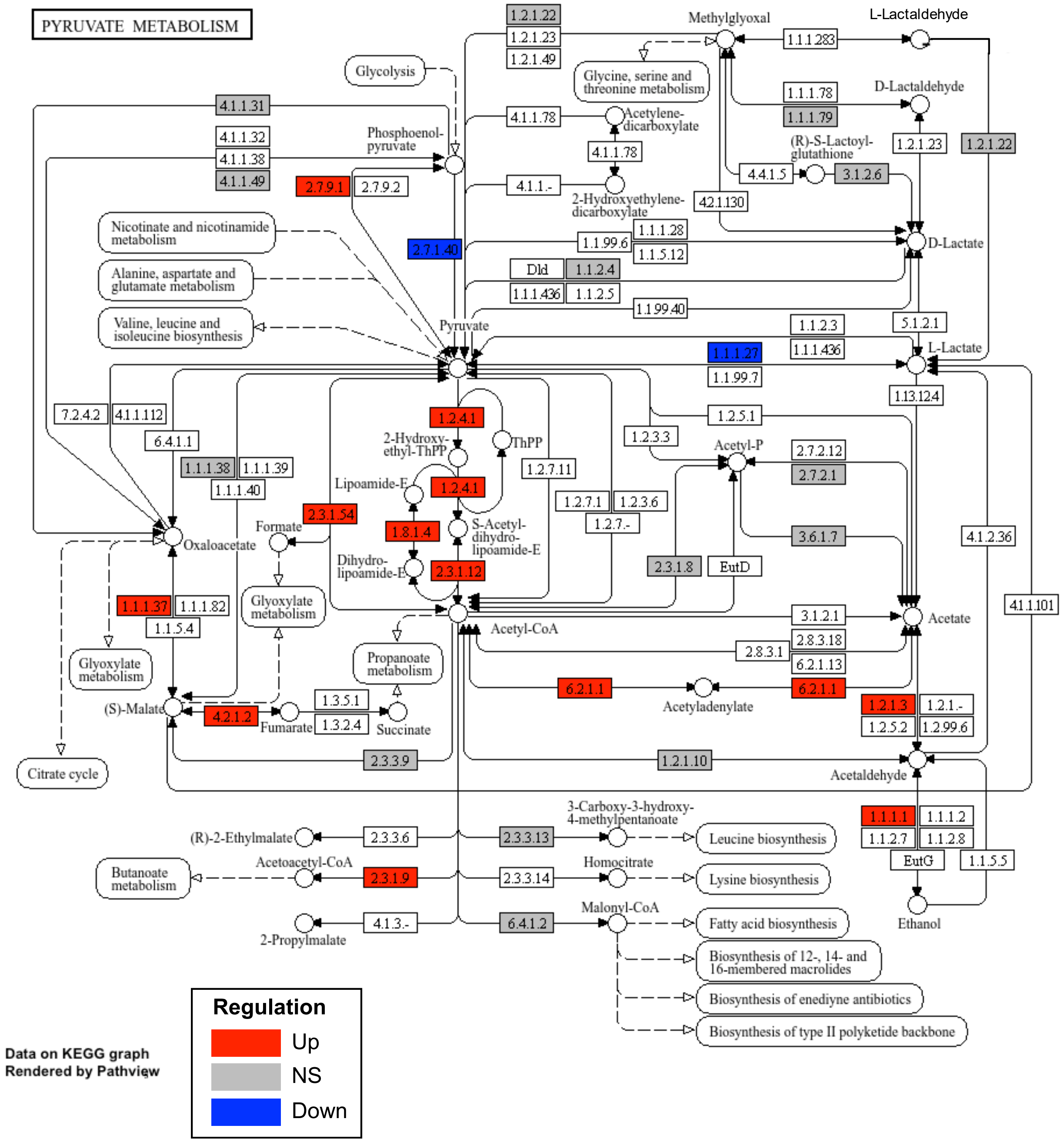


**Figure S4:** Differential expression of proteins involved in Pyruvate Metabolism of *B. coagulans* B-768 growing on xylose and glucose. “Up” regulation, adjusted p-value < 0.05 and log2 fold change > 1; “NS” = Not Significant” “Down” regulation, adjusted p-value < 0.05 and log2 fold change < -1.

### Differential Expression of Xylose vs. Glucose

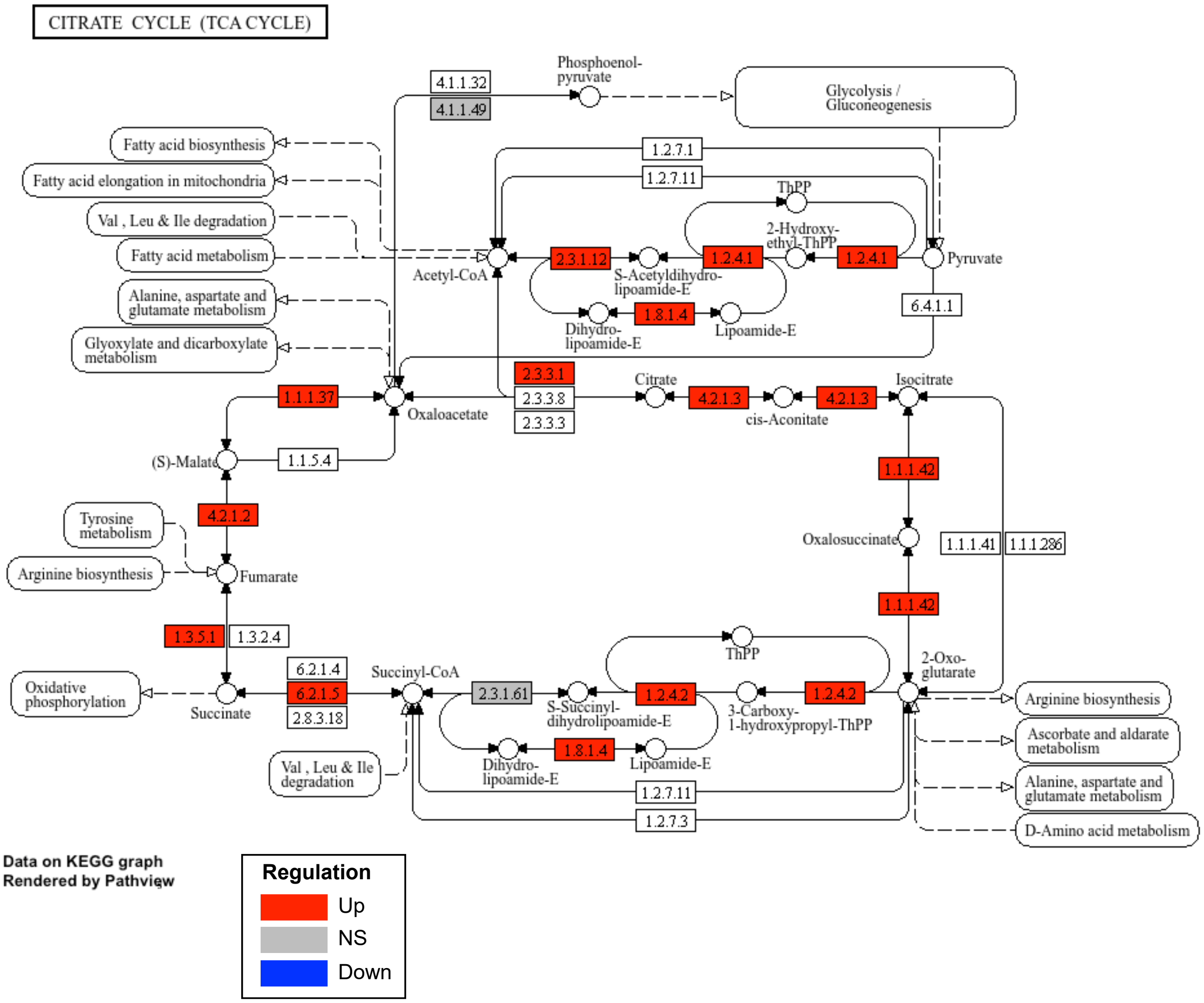


**Figure S5:** Differential expression of proteins involved in TCA cycle of *B. coagulans* B-768 growing on xylose and glucose. “Up” regulation, adjusted p-value < 0.05 and log2 fold change > 1; “NS” = Not Significant” “Down” regulation, adjusted p-value < 0.05 and log2 fold change < -1.
